# A ggplot-based single-gene viewer reveals insights into the translatome and other nucleotide-resolution omics data

**DOI:** 10.1101/2025.01.30.635743

**Authors:** Hsin-Yen Larry Wu, Isaiah D. Kaufman, Polly Yingshan Hsu

**Affiliations:** Department of Biochemistry & Molecular Biology, Michigan State University, East Lansing, MI 48824 USA

**Keywords:** ribosome profiling, visualization, periodicity, single-nucleotide resolution, translation

## Abstract

Seeing is believing. Visualizing Ribo-seq and other sequencing data within genes of interest is a powerful approach to studying gene expression, but its application is limited by a lack of robust tools. Here, we introduce *ggRibo*, a user-friendly R package for visualizing individual gene expression, integrating Ribo-seq, RNA-seq, and other genome-wide datasets with flexible scaling options. *ggRibo* visualizes 3-nucleotide periodicity, a hallmark of translating ribosomes, within a gene-structure context, including introns and untranslated regions, enabling the study of novel ORFs, translation of different isoforms, and mechanisms of translational regulation. *ggRibo* can plot multiple Ribo-seq/RNA-seq datasets from different conditions for comparison. It also contains functions for plotting single-transcript view, reading-frame decomposition, and RNA-seq coverage alone. Importantly, *ggRibo* supports the visualization of other omics datasets that could also be presented with single-nucleotide resolution, such as RNA degradome, transcription start sites, translation initiation sites, and epitranscriptomic modifications. We demonstrate its utility with examples of upstream ORFs, downstream ORFs, nested ORFs, and differential isoform translation in humans, *Arabidopsis*, tomato, and rice. We also provide examples of multi-omic comparisons that reveal insights that connect the transcriptome, translatome, and degradome. In summary, *ggRibo* is an advanced single-gene viewer that offers a valuable resource for studying gene expression regulation through its intuitive and flexible platform.

## INTRODUCTION

Visualization is emphasized in cell biology research to generate hypotheses and results. However, in omics studies, the visualization of sequencing data within genes of interest remains underutilized due to a lack of appropriate tools. Recent advancements in plotting tools in the R programming language now offer opportunities to develop specialized software packages for multi-omic data visualization.

Ribo-seq is a widely used technique for studying noncanonical open reading frames (ORFs) and mechanisms of mRNA translation at single-nucleotide resolution (Ingolia et al. 2009; Brar and Weissman 2015; Wu et al. 2024b). By optimizing RNase digestion to obtain ribosome-protected mRNA fragments (ribosome footprints), Ribo-seq captures the codon-by-codon, 3-nucleotide (3-nt) movement of ribosomes during translation, a feature known as 3-nt periodicity (Ingolia et al. 2009). During data analysis, specific single-nucleotide positions can be assigned to each ribosome footprint and color-coded by reading frame, enabling visualization of 3-nt periodicity along translated ORFs (Calviello et al. 2016; Hsu et al. 2016). Detecting 3-nt periodicity is crucial for identifying noncanonical ORFs and investigating the mechanism of mRNA translation (Guydosh and Green 2014; Chothani et al. 2023; Wu et al. 2024a).

While several Ribo-seq software packages generate aggregate plots to examine 3-nt periodicity at a global level (Ji et al. 2015; Calviello et al. 2016; Zhang et al. 2017; Xiao et al. 2018; Calviello et al. 2020; Choudhary et al. 2020; Harnett et al. 2021), visualizing 3-nt periodicity within individual genes remains challenging due to the complexity of gene structure and isoforms. Many studies simply present Ribo-seq data as coverage plots or single-color lines for genes of interest, disregarding the rich information offered by 3-nt periodicity. Others may depict periodicity within a single transcript isoform (Kiniry et al. 2019), assuming the selected isoform is the main expressed/translated isoform, thereby missing the opportunity to examine other annotated or unannotated isoforms. Although some studies display Ribo-seq in a genome browser, providing gene structure and isoform context, the reads are still shown in one color, preventing the intuitive visualization of 3-nt periodicity (Michel et al. 2018). Additionally, many available Ribo-seq visualization tools plot one sample at a time, making comparisons between samples challenging.

We previously developed *RiboPlotR* to visualize Ribo-seq reads for all annotated isoforms of individual genes within their gene structures, including 5′ and 3′ untranslated regions (UTRs) and introns. It displays both RNA-seq coverage and Ribo-seq periodicity by color-coding reads from each reading frame (Wu and Hsu 2021). Although *RiboPlotR* enables the visualization of isoform translation and many unannotated translation features, it has several limitations: 1) It only accepts a specific annotation format commonly used for plant gene annotation, where transcript IDs are assigned according to gene IDs plus a number. Therefore, *RiboPlotR* cannot plot data from human and many other non-plant species. 2) It only plots annotated ORFs and upstream ORFs (uORFs) in the 5′ UTR, not other unannotated ORFs, such as downstream ORFs (dORFs) in the 3′ UTR or novel small ORFs within annotated noncoding RNAs. 3) It only displays a maximum of two Ribo-seq samples per plot, preventing large-scale comparisons. 4) It only accepts a tabular input file for Ribo-seq and bam files for RNA-seq, but not other standard input files, such as bedGraph or bigWig. 5) It does not display DNA and amino acid sequences, which aid in identifying ORFs or specific sequence features. 6) It relies on base R plotting capabilities, producing figures that are less refined than those generated with *ggplot2* (Wickham 2016).

To enhance the flexibility and quality of Ribo-seq data visualization, here we introduce *ggRibo*, which extends the advantages of *RiboPlotR* while leveraging the capabilities of *ggplot2*. *ggRibo* aims to support broader species and input formats, visualize more types of unannotated ORFs, accommodate as many Ribo-seq/RNA-seq datasets as the computer’s RAM allows, display DNA and amino acid sequences when necessary, and, most importantly, integrate other sequencing data types with single-nucleotide resolution, where specific nucleotide positions convey sequence read information. Together, *ggRibo* enables parallel comparisons of multi-omic datasets, facilitating the study of gene regulation at multiple levels and revealing novel biological insights. We provide examples of using *ggRibo* to visualize different types of noncanonical ORFs in humans, *Arabidopsis*, rice, and tomato. Additionally, we present examples that demonstrate how *ggRibo* can be applied to gain an in-depth understanding of gene regulation using multi-omics data in Arabidopsis.

## RESULTS

### Design of *ggRibo*

**Figure 1A-C** illustrates how ribosome footprints can be represented by single-nucleotide positions (often the first nucleotide of the peptidyl site [P-site] within ribosomes) and color-coded by reading frames, enabling visualization of 3-nt periodicity along translated ORFs. P-site positions can be assigned using tools such as *RiboTaper* or *Ribo-seQC* (Calviello et al. 2016, 2019). Note that the P-sites for different ribosome footprint lengths may vary across organisms and organelles (**Figure 1A, D, E**) and depend on the footprinting conditions (Ingolia et al. 2012; Bazzini et al. 2014; Hsu et al. 2016; Chen et al. 2020; Wu et al. 2024a).

**Figure 1.**
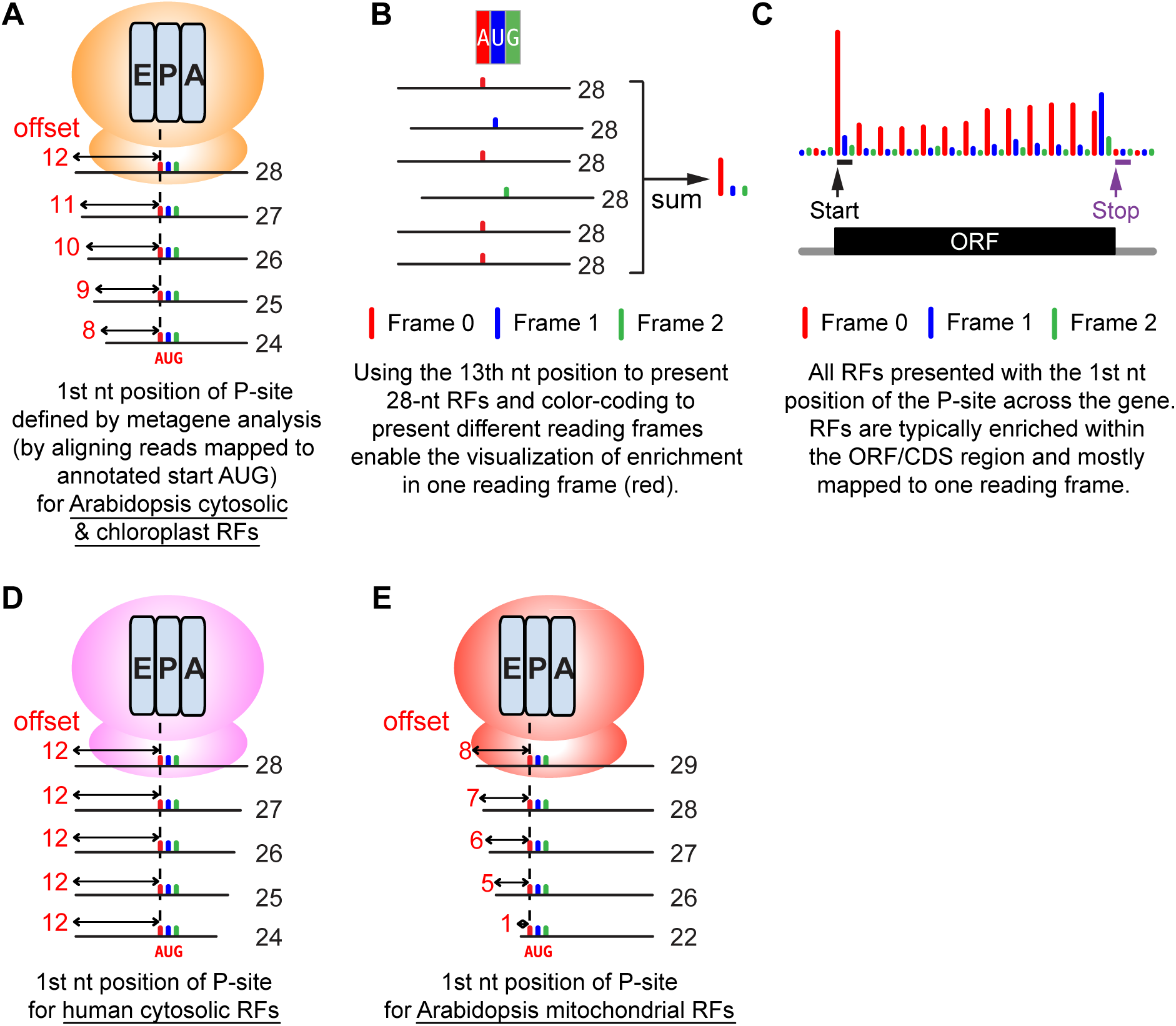
An illustration of how ribosome footprints (RFs) can be represented by the first nucleotide (nt) of the P-site and color-coded by reading frames to highlight 3-nt periodicity. (A) Metagene analysis by aligning all RFs mapped to the annotated start codon (AUG) can reveal the position of the P-site within ribosomes for each length of RFs. For *Arabidopsis* cytosolic and chloroplast RFs, the 13^th^ nt (offset by 12 nt from 5′) is typically the 1^st^ nt position of the P-site for a 28-nt RF. Similarly, the 12^th^ nt (offset by 11 nt from 5′) is the 1^st^ nt position of the P-site for a 27-nt RF. Note the P-site offsets could vary in different organisms and organelles (see panels D-E) (Ingolia et al. 2011; Hsu et al. 2016; Wu et al. 2024a). (B) An illustration showing how 28-nt RFs are mapped to a codon. The 28-nt RFs are presented by their 13^th^ nt position and color-coded by reading frame (annotated frame = frame 0: red, frame 1: blue, frame 2: green). Summing the reads mapped to each of the three nucleotides reveals the enrichment in frame 0 (red), which is expected in high-quality datasets. (C) Combining all RFs mapped to a particular gene produces a P-site plot, which visualizes the distribution of the RFs and colored 3-nt periodicity across the gene. At the codon prior to the stop codon, ribosome conformational change at translation termination causes RFs to have a different length and results in an enrichment of frame 1 (blue) (Wu et al. 2024a). (D) The P-site offsets for human ‘cytosolic’ RFs from induced pluripotent stem cells (iPSCs) and cardiomyocytes (Chen et al. 2020). Note that RNase digestion for different lengths of RFs occurs at the 3′ of the RFs in humans, compared to the 5′ of the RFs in *Arabidopsis* (see panel A). (E) The P-site offsets for *Arabidopsis* ‘mitochondrial’ RFs. The P-site offsets are shorter than the cytosolic and chloroplast RFs in *Arabidopsis* (see panel A).

To enhance the visual interpretation of periodic Ribo-seq reads, we developed *ggRibo* (https://github.com/hsinyenwu/ggRibo), a single-gene viewer that displays Ribo-seq periodicity alongside RNA-seq coverage in the context of the gene structure, including 5′/3′ UTRs and introns from multiple isoforms (**Figure 2A**). *ggRibo* supports all GTF and GFF annotation files compatible with the *GenomicRanges* package (Lawrence et al. 2013). *ggRibo* accepts common standard input files, such as bedGraph, bigWig, and bam. Built on the *ggplot2* framework, *ggRibo* generates publication-quality plots. To simplify interpretation, *ggRibo* displays genes from left to right, regardless of whether they are on the Watson or Crick strand. Ribo-seq reads are plotted by the first nucleotide of the P-site and color-coded by reading frames (**Figures 1** and **2A**). Together, these features facilitate the detection and the detailed visualization of unannotated ORFs and noncanonical translation features.

**Figure 2.**
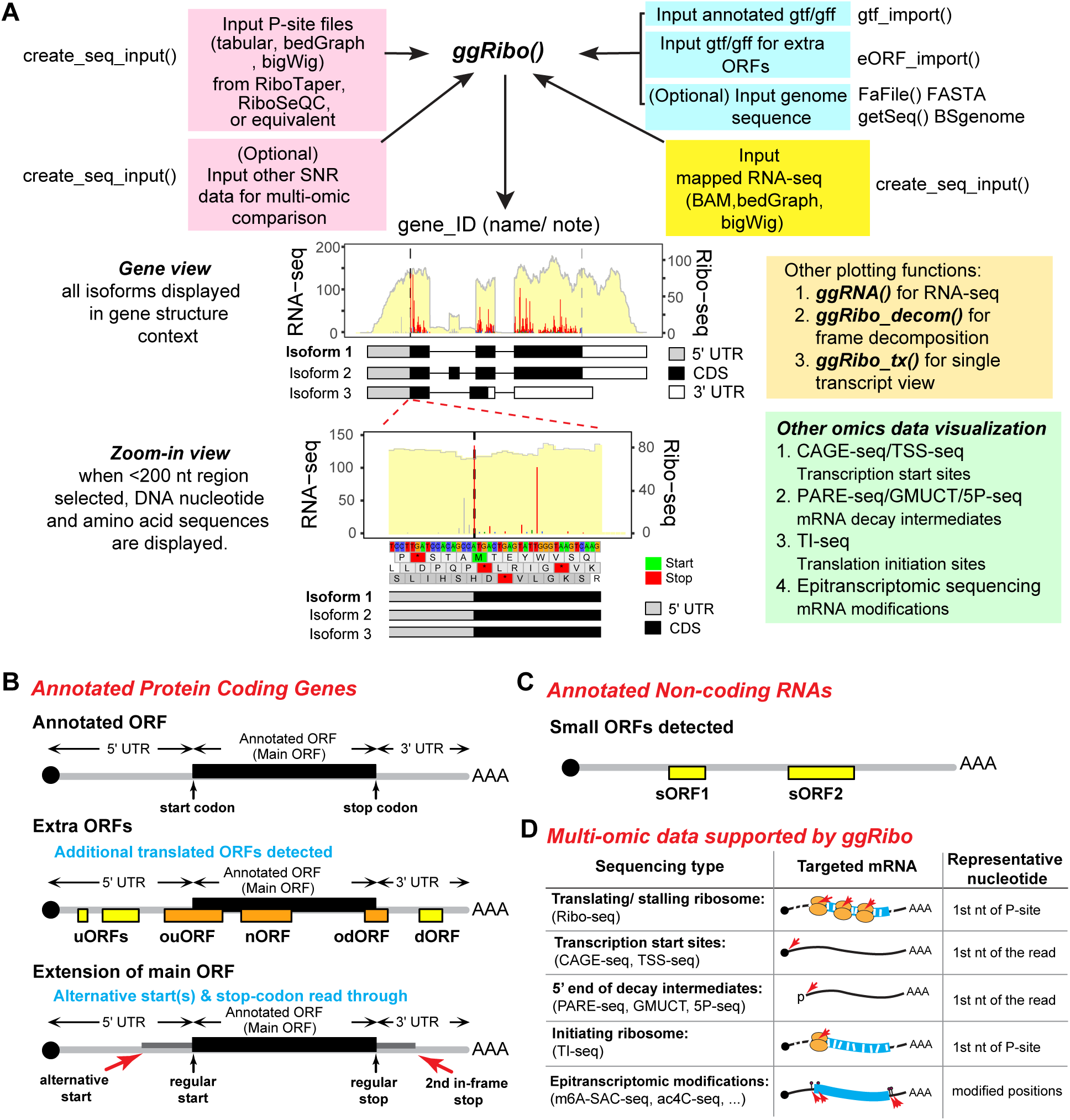
Overview of *ggRibo* and the ORF/data types supported. (A) Input/output and the functions of *ggRibo. ggRibo* requires Ribo-seq P-site information, genome annotation for annotated ORFs and (optional) extra ORFs, and mapped RNA-seq file(s). Other omics data presented with single-nucleotide resolution (SNR) can be input and visualized in parallel or independently. To visualize DNA and/or amino acid sequences, users can optionally provide the corresponding genome sequence file. Output plots can be in various scales, such as gene view or zoomed-in view. (B, C) ORF types can be visualized by *ggRibo*. Annotated genes are typically grouped into either coding (B) or noncoding (C), and *ggRibo* can plot ORFs in both gene types. For annotated protein-coding genes, typically only one ORF (main ORF) is annotated, while for annotated noncoding RNA genes, no ORF is annotated by default. By providing GTF/GFF files that include extra ORFs, users can visualize any ORF beyond those already annotated. ‘Extra ORFs’ may be translated using a frame different from the main ORF and/or separate from the main ORF, including upstream ORFs (uORFs) within 5′ UTR and downstream ORFs (dORFs) within 3′ UTR. Some of the uORFs and dORFs may partially overlap with the main ORF (ouORFs and odORFs). Some overlapping ORFs may be nested within the main ORF (nORF). In contrast, ‘Extension of main ORFs’ uses the same reading frame as the main ORF and extends from 5′ or 3′. The fExtend and tExtend parameters allow extending the main ORF frames into the 5′ UTR and 3′ UTR, respectively. This enables investigating translation from alternative AUG or non-AUG start codons in the 5′ UTR or stop-codon readthrough events in the 3′ UTR. For annotated noncoding RNAs, some small ORFs (sORFs) may be translated. (D) Example omics data types with single-nucleotide resolution that can be plotted by *ggRibo*.

Notably, *ggRibo* visualizes both annotated ORFs and extra ORFs (called “eORFs” in the *ggRibo* package) (**Figure 2B**). It can also extend the main ORF reading frame into the 5′ UTR and 3′ UTR, allowing the visualization of alternative start sites (AUG or non-AUG) and stop codon readthrough, respectively (**Figure 2B**). Additionally, *ggRibo* can plot Ribo-seq reads on annotated noncoding RNAs to visually inspect whether a potential small ORF is translated with strong 3-nt periodicity (**Figure 2C**). A summary of *ggRibo*’s major improvements over its prototype, *RiboPlotR*, is described in **Supplemental Figure S1**.

*ggRibo* is a user-friendly R package that accepts a wide range of standard input files, is applicable to any species with a genome annotation, and offers improved flexibility for customization and easy automation compared to other existing tools, like *GWIPS-viz*, *Trips-Viz,* and *RiboCrypt* (Michel et al. 2018; Kiniry et al. 2019) (https://ribocrypt.org/). *GWIPS-viz* and *Trips-Viz* are online genome browsers that display a collection of data for specific species and genome annotations, while *RiboCrypt* offers both online and user-installed genome browsers when used in conjunction with the *ORFik* package in R (Tjeldnes et al. 2021). In contrast, *ggRibo* can be easily integrated into any pipeline by offering simple functions for loading and visualizing data. Importantly, *ggRibo* provides an efficient presentation of gene structure, isoform expression/translation, and color-coded reading frame information simultaneously in high-quality figures (**Supplemental Figure S2A**). A single transcript view option is also available using *ggRibo_tx* (**Supplemental Figure S3A**). In contrast to *ggRibo*, *GWIPS-viz* plots Ribo-seq reads in a single color, which hinders the intuitive interpretation of reading frames (**Supplemental Figure S2B**). Although *RiboCrypt* offers color-coded reading frames in ‘gene’ view, the reading frame is assigned based on the genomic sequence, and therefore, the colors change as the exon sequences shift from one reading frame to another (**Supplemental Figure S2C**). The change in color due to a reading frame shift makes it challenging to identify the ORF visually, which is crucial for studying multiple or overlapping ORFs in one transcript (see below; **Figures 4E, F,** and **6A**). Notably, *ggRibo* displays a consistent color for an ORF as the reading frame is assigned based on the coding sequence (**Supplemental Figure S2A**). Taken together, compared to existing tools, *ggRibo* offers intuitive visualization and superior flexibility for customization.

In addition to Ribo-seq/RNA-seq, *ggRibo* supports the visualization of other sequencing data that are presented with single-nucleotide resolution (**Figure 2D**). Examples include CAGE-seq and TSS-seq for studying transcription start sites (Wakaguri et al. 2008; Valen et al. 2009); PARE-seq, GMUCT, and 5P-seq for studying mRNA decay intermediates (German et al. 2008; Gregory et al. 2008; Pelechano et al. 2015); TI-seq for studying translation initiation sites (Ingolia et al. 2011; Lee et al. 2012; Willems et al. 2017; Li and Liu 2020), and m6A-SAC-seq and ac4C-seq for studying epitranscriptomic modifications (Sas-Chen et al. 2020; Hu et al. 2022).

*ggRibo* package extends its capabilities beyond the *ggRibo* function by including three specialized tools: *ggRNA*, *ggRibo_decom*, and *ggRibo_tx*; each tailored for specific visualization tasks.

#### ggRNA: RNA-seq Data Visualization

*ggRNA* plots RNA-seq data alone, when only RNA-seq coverage is needed (**Supplemental Figure S4A**). Like the *ggRibo* function, it supports multiple samples and accommodates both paired-end and single-end RNA-seq data, offering flexibility for various experimental designs.

#### ggRibo_decom: Reading Frame Decomposition

*ggRibo_decom* plots Ribo-seq reads from different reading frames separately for analyzing frame-specific distributions (**Supplemental Figure S4B**). One dataset is processed at a time for in-depth examination.

#### ggRibo_tx: Single-Transcript Visualization

*ggRibo_tx* visualizes RNA-seq and Ribo-seq data in the context of mature RNA for a selected transcript isoform (**Supplemental Figure S4C**), which is useful for genes with long or numerous introns. To prevent misinterpretation from incorrect isoforms, users should first identify expressed isoforms using *ggRibo* or *ggRNA* before employing *ggRibo_tx*. Similar to the *ggRibo* function, *ggRibo_tx* displays both RNA-seq and Ribo-seq data from multiple samples in a single plot.

#### Applications and Further Information

Below, we present examples demonstrating how *ggRibo* and its specialized functions can be used to explore unannotated ORFs and noncanonical translation features within the translatome, as well as how it can integrate multi-omics datasets to enhance our understanding of gene regulation. A tutorial detailing the setup, requirements, and example code of the *ggRibo* package is available on the *ggRibo* GitHub site.

### Visualizing Translated ORFs with Human and Rice Data

To demonstrate *ggRibo*’s basic usage, we first provide an example in the human *MRPL11* gene (**Figure 3**), which contains a uORF in the 5′ UTR that represses downstream main ORF translation (Calvo et al. 2009). **Figure 3A** presents the Ribo-seq and RNA-seq data of *MRPL11* in human embryonic stem cells (Chothani et al. 2022), displaying all annotated RNA isoforms, with the uORF highlighted in yellow in the gene models. **Figure 3B** adds an amino acid track with start and stop codon positions highlighted. **Figure 3C** zooms in on the *MRPL11* uORF, displaying both DNA nucleotide and amino acid sequence tracks.

**Figure 3.**
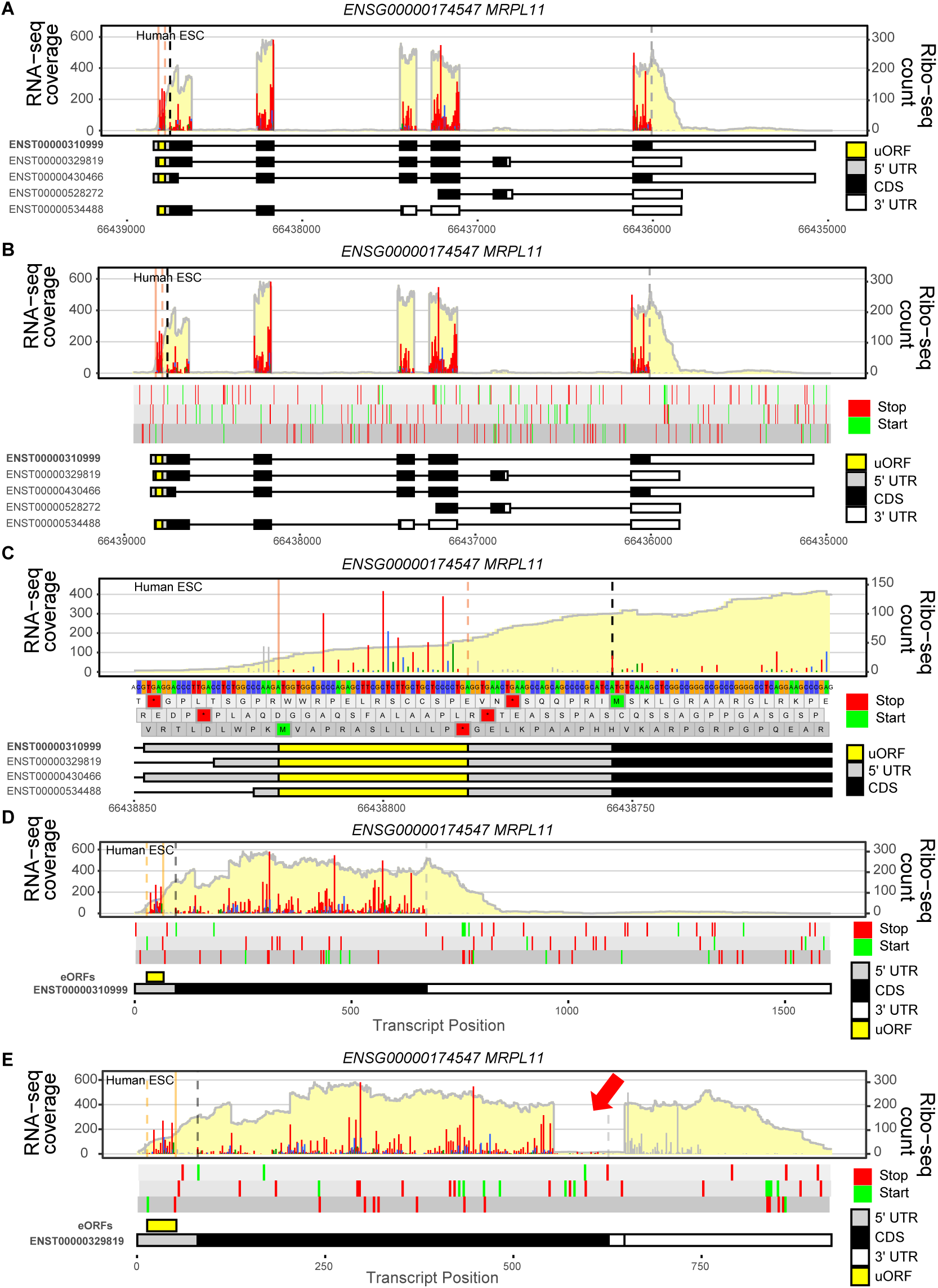
Examples of *ggRibo* and *ggRibo_tx* plots for the human *MRPL11* gene. Different display options using *ggRibo* to present RNA-seq and Ribo-seq profiles of the human nuclear-encoded *MRPL11* gene. The human embryonic stem cell (ESC) Ribo-seq and RNA-seq data are from (Chothani et al. 2022). RNA-seq coverage is shown with a light-yellow background. Ribo-seq reads are presented with their first nt of the P-site, and they are color-coded in red, blue, and green to indicate they map to reading frame 0 (expected), 1, and 2, respectively. *MRPL11* possesses 5 annotated transcript isoforms, and both a uORF and the main ORF are translated. The user can specify which transcript isoform to focus on and assign the reading frame according to its gene structure; here, ENST00000310999 is selected (bolded). Within the gene models, black boxes indicate the coding sequence (CDS) (i.e., the annotated/main ORF), yellow box(es) indicate the uORF region, and gray and white boxes indicate 5′ UTR and 3′ UTR, respectively. Black and gray vertical dashed lines represent the start and stop codons for the annotated mORF, respectively. Orange solid and dashed vertical lines represent start and stop codons for the uORF, respectively. Reads outside of the ORF range are shown in grey. Genomic or transcript coordinates on the chromosomes are indicated below the gene models. (A) Only Ribo-seq and RNA-seq profiles are presented on top of the annotated gene models. (B) An additional DNA sequence track was added, but only the start and stop codons are displayed due to the large span (>201 nt). (C) The view is further zoomed in on the uORF region, showing both the DNA nucleotide and amino acid sequence tracks in detail (when length <201 nt). Note that the line width of the Ribo-seq reads could be modified by the “*ribo_linewidth*” parameter, and the sequence color may be modified by the “*nucleotide_color_scheme*” parameter (D) Single-transcript plot by *ggRibo_tx* to show the RNA-seq and Ribo-seq reads that map to the main expressed isoform ENST00000310999. (E) Similar to (D), but for a very lowly expressed isoform ENST00000329819 in this dataset (See **Figure 3A**). The arrow highlights exon 5, which contains a small number of Ribo-seq reads with 3-nt periodicity.

To focus on the expressed isoforms, we next use *ggRibo_t*x to visualize *MRPL11*. **Figure 3A** suggests that the main expressed isoform is ENST00000310999. Therefore, we plot this isoform with *ggRibo_tx* in **Figure 3D**. To demonstrate the importance of visualizing the data within the gene context, we also plot the isoform ENST00000329819, which is expressed and translated at low levels relative to ENST00000310999, as shown by the small number of periodic reads in exon 5 of ENST00000329819 (**Figure 3E**).

Next, we examine the human immune-related gene *IFITM3* (*Interferon-Induced Transmembrane Protein 3*), which contains isoforms with and without an annotated coding sequence, using human brain data (Chothani et al. 2022). *ggRibo* plot reveals that ENST00000399808 is the major expressed and translated isoform (**Supplemental Figure S5A**: a gene view displaying all RNA isoforms, **Supplemental Figure S5B**: a zoom-in view of a subset of isoforms). This example again highlights how visualizing RNA-seq and Ribo-seq reads in the context of gene structure helps identify expressed and translated isoforms.

Similarly, we provide an example in rice to visualize a uORF encoding a conserved peptide in the *AdoMetDC* gene (Franceschetti et al. 2001; Hayden and Jorgensen 2007) using *ggRibo* and *ggRibo_tx plots* (**Supplemental Figure S6**) with the data reported in (Yang et al. 2021). The striking difference in Ribo-seq levels between the uORF and the main ORF suggests that the uORF almost completely suppresses the translation of the main ORF (**Supplemental Figure S6**).

### Visualizing Translation of Unannotated ORFs and within Noncoding RNAs in Arabidopsis and Tomato

We next present four examples to showcase how *ggRibo* facilitates the identification and visualization of novel isoforms, unannotated ORFs, as well as translation detected within noncoding RNAs using *Arabidopsis* and tomato data (Wu et al. 2019, 2024a).

#### First Example: Translation of an unannotated isoform

*PAE7* encodes a pectin acetylesterase and has two annotated RNA isoforms in Arabidopsis. RNA-seq coverage indicates that both isoforms are expressed, and the Ribo-seq periodicity supports that both isoforms are translated (**Figure 4A**). Additionally, cross-referencing the RNA-seq profile with gene models reveals that exon 14 of isoform 2 is longer than the annotated range (**Figure 4A**, blue dashed box). This finding further demonstrates how gene-structure-based visualization aids in interpreting isoform translation and refining gene annotations.

**Figure 4.**
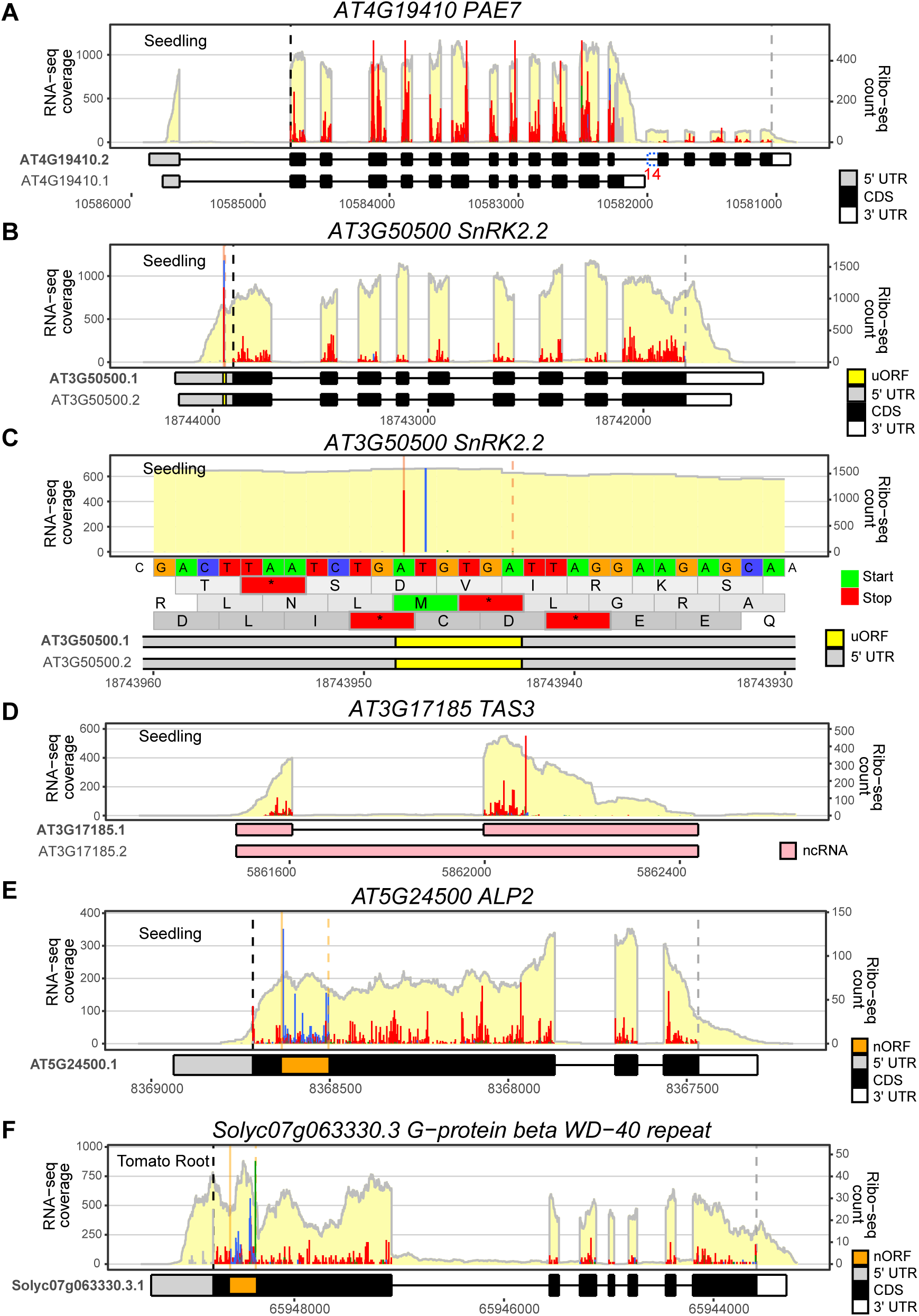
Examples of *ggRibo* plots with *Arabidopsis* and tomato genes. The *Arabidopsis* seedling Ribo-seq and RNA-seq data are from (Wu et al. 2024a). The tomato Ribo-seq and RNA-seq data are from (Wu et al. 2019). Data presentations are as described in Figure 3 legend. (A) Gene view *of PAE7* shows that both isoforms are translated. Note that the RNA-seq coverage reveals that the 14^th^ exon (highlighted) is longer than the annotated exon range for this isoform. The inferred exon 14 extension is shown with a blue dashed box (added with Adobe Illustrator). (B-C) A gene view and zoom-in view show the strong minimum uORF (i.e., AUG-stop) in *SnRK2.2*. In (C), the Ribo-seq reads precisely map to the ‘A’ and ‘T’ of the start codon. Orange solid and dashed vertical lines represent the translation start and stop, respectively, for the uORF. (D) A small ORF is translated within *TAS3*, an annotated noncoding RNA (ncRNA). Robust 3-nt periodicity is detected in the 50-aa small ORF region. (E) A nested ORF (nORF) using the blue reading frame is detected in the *Arabidopsis APL2* gene. The nested ORF range is highlighted with an orange box in the gene model. (F) A nested ORF using the blue reading frame is detected in a tomato G-protein-coupled receptor beta subunit gene. The nested ORF range is highlighted with an orange box in the gene model.

#### Second Example: Translated minimum uORF (AUG-STOP)

*SnRK2.2* encodes a key regulator of plant abiotic stress responses, and previous research identified a minimal uORF (AUG-STOP, the shortest possible ORF) in this gene (Wu et al. 2024a). Here, we plot the entire *SnRK2.2* gene (**Figure 4B**) and a zoomed-in view with the DNA nucleotide and amino acid sequence tracks centered on the uORF (**Figure 4C**). The Ribo-seq reads for this minimal uORF are predominantly mapped to the first two nucleotides, A (frame 0, red) and U (frame 1, blue) (**Figure 4C**). This read distribution, which is uniquely associated with minimal uORFs, results from the altered ribosome footprint length at the codon preceding the stop codon (Wu et al. 2024a).

#### Third Example: Translation of noncoding RNA

*TAS3* is an annotated noncoding RNA, encoding the primary transcript of trans-acting short-interfering RNA (*tasiRNA*). A small ORF is known to be translated within *TAS3* primary transcript, a process that regulates tasiRNA biogenesis (Hsu et al. 2016; Li et al. 2016; Bazin et al. 2017; Hsu and Benfey 2018; Iwakawa et al. 2021; Wu et al. 2024a). Using *ggRibo*, we can observe that Ribo-seq reads exhibit strong 3-nt periodicity at the region corresponding to this small ORF (**Figures 4D** and **Supplemental Figure S7**: zoom-in view showing the sequences near the start and stop codons). Further analysis using *ggRibo_decom*, which plots three-frame reads separately, confirms that the Ribo-seq reads are predominantly enriched in frame 0 (**Supplemental Figure S8**).

Notably, for noncoding RNAs, *ggRibo* assigns the reading frame from the first nucleotide of the annotated RNA sequence, rather than from the start of an ORF, as it does for coding RNAs. As a result, a translated ORF in a noncoding RNA may use one of the reading frames in red, blue, or green. **Supplemental Figure S9** shows an example of a small ORF enriched in the ‘blue’ reading frame from an annotated long noncoding RNA (lncRNA) in Arabidopsis.

#### Fourth Example: ORFs within ORFs, visualizing overlapping/nested translation events

We next show two examples of nested ORFs, in which two ORFs using different reading frames are both translated, and the smaller ORF is embedded within the longer one. While this phenomenon was discovered decades ago in bacterial phage φX174 (Barrell et al. 1976; Sanger et al. 1977), it is challenging to detect such nested ORFs in eukaryotes (Wright et al. 2022). Here, we attempt to visualize such events using high-coverage super-resolution Ribo-seq data with *ggRibo*. In *Arabidopsis ALP2* gene, a nested ORF using the blue reading frame (corresponding to the orange box in the transcript model) is clearly visible (**Figure 4E**). Similarly, in tomato *Solyc07g063330.3* gene, using data from (Wu et al. 2019), a nested ORF using the blue reading frame is also clearly visible (**Figure 4F**). Strong 3-nt periodicity in the nested ORF regions suggests that they are actively translated (**Figure 4E, F**). Frame decomposition using *ggRibo_decom* further supports these findings (**Supplemental Figure S10A, B**). The *ggRibo_decom* function is particularly useful for illustrating such frame enrichment for complex translation events.

### Comparing Differential Translated Isoforms Across Datasets using *ggRibo*

*ggRibo* can visualize multiple Ribo-seq/RNA-seq datasets to compare differential translated isoforms and/or translation efficiency of genes of interest. Here, we present an example of tissue-specific isoform translation in *Arabidopsis BCA4* (*β-Carbonic Anhydrase 4*), comparing the root, shoot, and whole seedling. *BCA4* plays a crucial role in gas exchange between plant leaves and the atmosphere, catalyzing the conversion of CO₂ to HCO₃⁻ and contributing to the stomatal CO₂ response (Hu et al. 2010, 2015). *BCA4* has three annotated isoforms; *ggRibo* plot suggests that isoforms 2 and 3 are expressed and translated in the tissues examined (**Supplemental Figure S11**). Interestingly, while isoform 2 is highly expressed and translated in the shoot and whole seedling, it is nearly undetectable in the root. Instead, the root mainly expresses the isoform 3 (**Supplemental Figure S11**). Recent work has shown that isoform 2 protein (the longer isoform) localizes to the plasma membrane, while isoform 3 protein (the shorter isoform) localizes to the cytosol (Weerasooriya et al. 2024). This example highlights *ggRibo*’s capacity to compare multiple datasets, enabling investigations into differential translation or isoform-specific translation across tissues/ conditions and providing valuable insights for future functional characterization.

### Multi-omic Data Comparisons Using *ggRibo*

Besides visualizing Ribo-seq/RNA-seq profiles, *ggRibo* can integrate other omics datasets that offer single-nucleotide resolution (See **Figure 2D**). To illustrate its utility in multi-omic data comparison, we provide three examples below to showcase *ggRibo*’s capacity to reveal novel biological insights for complex gene expression regulation. Note that other single-nucleotide-resolution data can be displayed either with a single color (example in **Figure 5A**) or with three colors to view the reading frame information (examples in **Figures 5B, 5C**) by setting the ‘sample_color’ parameter in the *ggRibo* function. Other omics datasets should be formatted similarly to Ribo-seq and input using the *create_seq_input* function.

**Figure 5.**
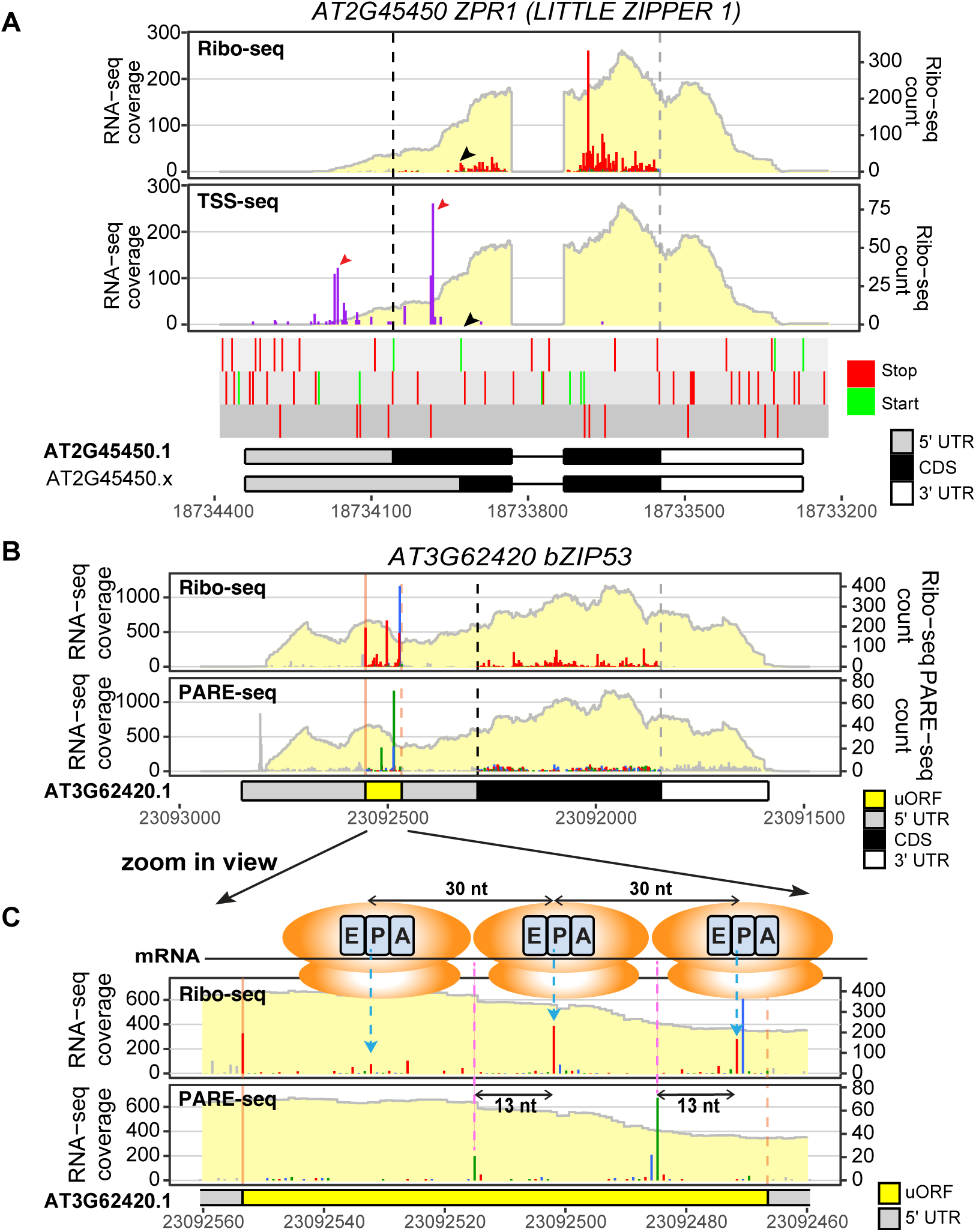
Multi-omic comparisons reveal novel insights connecting the transcriptome, translatome, and degradome. (A) A gene view showing Ribo-seq/RNA-seq (Wu et al. 2024a) and TSS-seq (Nielsen et al. 2019) profiles for AT2G45450 *LITTLE ZIPPER 1 (ZPR1)*. The Ribo-seq panel confirmed the translation of *ZPR1*, but the translation does not initiate at the annotated start codon (black dashed line), rather, reads mostly begin at the next in-frame AUG (black arrowhead in the Ribo-seq panel and start/stop codon track). TSS-seq (purple lines) reveals two major transcription start site (TSS) clusters (red arrowheads): one upstream and one downstream of the annotated start codon. The downstream TSS cluster will cause the translation to start at the second AUG and result in the 93-aa peptide. The color for TSS-seq data can be modified using the *sample_color* parameter in *ggRibo.* Ribo-seq and RNA seq data are reported in (Wu et al. 2024a), and TSS-seq data are reported in (Nagarajan et al. 2019). (B) A gene view showing Ribo-seq and PARE-seq in parallel for *bZIP53* gene; RNA-seq coverage is shown in a light-yellow background for each panel. The PARE-seq are also color-coded to compare the reading frame relationship with the Ribo-seq data. (C) A zoomed-in view of the conserved peptide uORF region. Accumulation of Ribo-seq reads at a 30-nt interval (highlighted by blue dashed arrows) upstream of the uORF stop codon (orange dashed line) suggests that multiple ribosomes stack in this region. Accumulation of the PARE-seq reads (highlighted by pink dashed lines) occurs at 13 nt upstream of the corresponding Ribo-seq peaks, supporting co-translational mRNA decay in this region. The shift between the red/blue reading frames prior to the stop codon in Ribo-seq is also observed in PARE-seq (blue/green). Ribosomes, blue dashed arrows, and pink dashed lines were added in Adobe Illustrator.

#### *First Example:* transcriptional start sites of *LITTLE ZIPPER 1* affect its translation

LITTLE ZIPPERS (ZPR1 to 4) are the first microproteins studied in *Arabidopsis* (Wenkel et al. 2007; Kim et al. 2008). Microproteins are single-domain proteins that usually bind to their multi-domain homologs and act as dominant negative regulators (Eguen et al. 2015; Bhati et al. 2021). While ZPR1 was predicted to be a 93-aa peptide (Wenkel et al. 2007), it is annotated as 136 aa in both TAIR10 and Araport11 annotations. To verify its actual translated peptide length, we examined *ZPR1* Ribo-seq profile using *ggRibo*. We found that most Ribo-seq reads for *ZPR1* locate downstream of the annotated AUG start (**Figure 5A: Ribo-seq panel**). A closer inspection revealed that the Ribo-seq reads mostly start at the next in-frame AUG, i.e., from the ORF in the *AT2G45450.x* isoform we defined (**Figure 5A**: black arrowheads in Ribo-seq panel and the start/stop codon track), supporting that *ZPR1* is predominantly translated into a 93 aa peptide.

Why does *ZPR1* primarily use the second AUG? Our recent study revealed that transcription start sites in *Arabidopsis* can be highly variable, affecting ORF translation initiation sites (Wu and Hsu 2024). Accordingly, some transcripts may begin transcription downstream of the annotated start codon (AUG), thereby excluding the first AUG from the mature mRNA. To test this possibility, we examined available TSS-seq data, which identify transcription start sites (TSSs) (Nielsen et al. 2019), and compared them with the Ribo-seq data and the gene model (**Figure 5A**). Consistent with our hypothesis, the TSS-seq signals for *ZPR1* occur primarily downstream of the annotated start codon, explaining why *ZPR1* translation preferentially initiates at the second AUG (**Figure 5A: TSS-seq panel**). This demonstrates how *ggRibo*’s capacity to integrate multi-omic datasets facilitates a deeper understanding of gene regulation.

#### *Second Example:* co-translational mRNA decay observed in a conserved peptide uORF

Co-translational mRNA decay is a process in which mRNA is degraded while still being translated or occupied by ribosomes. It is often detected at conserved peptide uORFs, where significant ribosome stalling occurs upstream of the stop codon (Hou et al. 2016; Yu et al. 2016; Guo et al. 2023). Here, we present an example in the *bZIP53* transcription factor gene, comparing Ribo-seq/RNA-seq profiles with PARE-seq data (Nagarajan et al. 2019) to evaluate potential mRNA decay associated with ribosome stalling (**Figure 5B**). PARE-seq detects the 5′-mono-phosphate end, which is a common mRNA decay intermediate (German et al. 2008). As noted in previous studies, Ribo-seq and PARE-seq suggest that ribosomes stall at approximately 30-nucleotide intervals (the estimated width of a ribosome footprint) from the end of the uORF (**Figure 5C**). Meanwhile, PARE-seq data display corresponding peaks 13 nucleotides upstream of the Ribo-seq peaks (**Figure 5C**). This pattern aligns with the expectations for co-translational decay, where ribosomes act as a protective barrier against nucleolytic degradation, causing mRNA decay to stop at the 5′ boundary of the ribosome (Hou et al. 2016; Yu et al. 2016). Remarkably, the shift between red and blue reading frames in Ribo-seq data prior to the stop codon (**Figure 5C, Ribo panel**) (Wu et al. 2024a) is also reflected in PARE-seq data (blue/green) (**Figure 5C, PARE panel**). Thus, visualization of multi-omic data using *ggRibo* allows in-depth analysis of gene regulatory mechanisms.

#### *Third Example:* a downstream ORF (dORF) that potentially overlaps with the main ORF

We previously reported that *AT1G49980*, which encodes a DNA/RNA polymerase family protein, harbors a strongly translated dORF in the 3′ UTR, using an ORF discovery tool, *RiboTaper* (Calviello et al. 2016; Wu et al. 2024a). *RiboTaper* classified this dORF as an overlapping dORF in *AT1G49980.1* isoform, overlapping with the stop codon of the main ORF (see gene models in **Figure 6A**). Plotting *AT1G49980* by *ggRibo* confirmed a highly translated ORF within the annotated 3′ UTR (**Figure 6A, Ribo-seq panel**: green reading frame). However, the higher RNA coverage in the dORF region compared to the main ORF region suggests that this dORF arises from a separate, unannotated gene or transcript. To test this possibility, we examined transcription start sites reported in TSS-seq data (Nielsen et al. 2019) by *ggRibo*. We found two distinct TSS clusters: one near the annotated transcription start site and the other upstream of the dORF (**Figure 6A-B, TSS-seq panel**: purple lines). This pattern suggests the presence of an unannotated gene/transcript downstream of *AT1G49980*, which is supported by an independent study (Thieffry et al. 2020). Therefore, this apparent dORF is more likely a novel ORF from an unannotated gene, rather than a dORF of *AT1G49980*.

**Figure 6.**
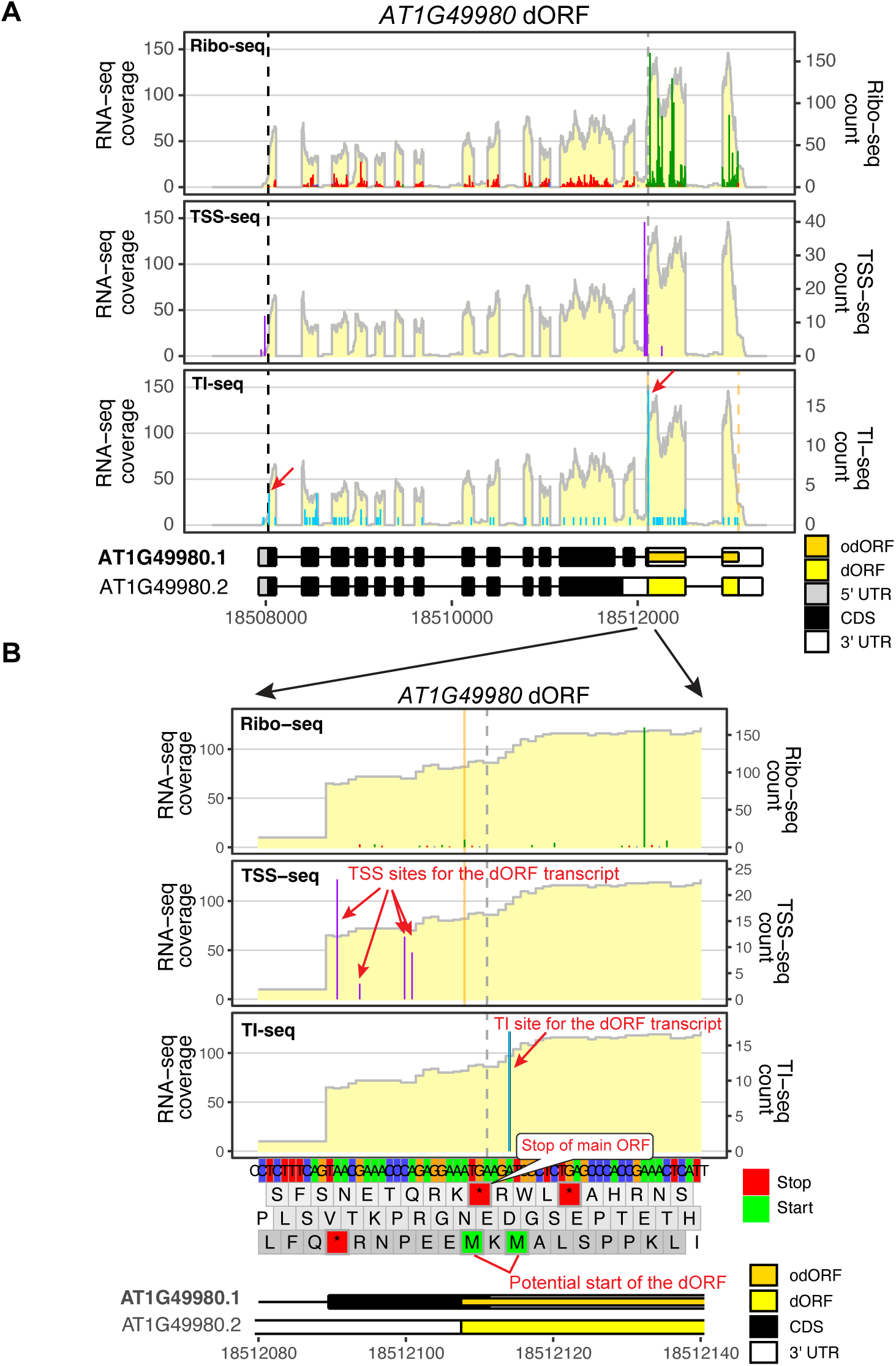
Multi-omic comparison reveals the nature of a translated “dORF”. Ribo-seq and RNA seq data are reported in (Wu et al. 2024a), TSS-seq data are reported in (Nielsen et al. 2019), and TI-seq data are reported in (Willems et al. 2017). (A) A gene view showing Ribo-seq, TSS-seq, and TI-seq in parallel for *AT1G49980*; RNA-seq coverage is shown in the background for each panel. Ribo-seq panel confirms a potential dORF translated in the 3′ UTR using frame 2 (green), which was previously identified by *RiboTaper* (Calviello et al. 2016). TSS-seq panel reveals 2 major TSS clusters (purple lines); one corresponds to the annotated transcription start site for *AT1G49980*, and the other upstream of this dORF. The existence of a transcription start site upstream of the dORF supports this dORF arising from an unannotated gene/transcript, which has higher RNA levels than *AT1G49980*. (B) A zoomed-in view focusing on the 5′ of the dORF using the same data above. The grey vertical dashed line indicates the stop codon of the main ORF, and the orange solid line indicates the *RiboTaper*-predicted start codon for the dORF, which corresponds to the first of the two adjacent ATG starts (indicated by two methionine amino acids). The TI-seq data reveals that this dORF exclusively uses the second start codon, and therefore, this ORF does not overlap with the main ORF stop codon. Note that *RiboTaper* predicted this dORF to overlap with the main ORF of isoform 1 but not the main ORF of isoform 2. Therefore, in isoform 1, this dORF is considered an overlapping dORF (colored orange in the isoform 1 gene model), while in isoform 2, it is considered a distinct dORF (colored yellow in the isoform 2 gene model). The red arrows and comments were added in Adobe Illustrator.

Further analysis of the dORF’s DNA sequence reveals 2 potential ATG start codons: one overlapping with the main ORF’s stop codon and the other located 2 codons downstream (**Figure 6B**, bottom panel). To determine which start codon is used by this dORF, we examined available TI-seq data (Willems et al. 2017), a technique similar to Ribo-seq but enriching for initiating ribosomes, which assists in the identification of translation initiation sites (Ingolia et al. 2011; Lee et al. 2012; Willems et al. 2017; Li and Liu 2020). The TI-seq data show 2 major peaks: the first one near the annotated start codon of *AT1G49980* and the second one at the dORF region (arrows in **Figure 6A, TI-seq panel**). Zooming in on the 5′ region of the dORF revealed that this dORF exclusively uses the second ATG (**Figure 6B, TI-seq panel**), indicating that this translated dORF does not overlap with the *AT1G49980* main ORF. This example again highlights *ggRibo*’s capacity to integrate multi-omic data with unparalleled resolution, which helps reveal novel gene regulatory mechanisms for future studies.

## DISCUSSION

In omics studies, the visualization of sequencing data within genes of interest remains underutilized due to a lack of appropriate tools. To address this gap, we developed *ggRibo*, which enables in-depth exploration of translational landscapes under various experimental conditions and facilitates multi-omics comparisons, providing deeper insights into translational regulation and different levels of gene regulation.

*ggRib*o and its associated plotting functions offer several advantages over web-based tools: (1) Flexible Visualization: Users can input and visualize RNA-seq, Ribo-seq (which exhibits 3-nt periodicity), and any other single-nucleotide resolution data together for genes of interest with flexible scaling. Other single-nucleotide resolution sequencing data and RNA-seq data can also be plotted alone, independent of Ribo-seq. (2) Batch Plotting: Users can plot hundreds of genes at once to visualize their genes of interest. For example, one could quickly compare uORF translation under different conditions across hundreds of uORF-containing genes. The same task might take days using genome or transcript browsers, which require manual copying and pasting for each gene. (3) Publication-Ready Output: The *ggRibo* output can be saved as PDF, SVG, PNG, or any other file format compatible with *ggplot2*’s ggsave function for easy data management and editing of figures for publication. (4) Custom ORF Input: Users can input a GTF file to include genomic coordinates of additional ORFs (such as uORFs and dORFs) to facilitate the visualization of noncanonical ORFs. (5) Isoform Confirmation: The gene view in *ggRibo* allows users to identify translated isoforms, whether annotated or unannotated. Relying solely on annotated isoforms in single-transcript viewers might overlook translation occurring on unannotated transcripts. (6) Availability: Online tools may be restrained by server capacity and only maintained as funding permits, but users can always use R and *ggRibo*.

In summary, with its flexibility and simple workflow, *ggRibo* is a powerful tool for multi-omics analysis. It helps researchers efficiently visualize individual gene expression from diverse sequencing data, produce publication-quality plots, synthesize hypotheses for future studies, and develop a more comprehensive understanding of gene regulation.

## METHODS

The *ggRibo* package provides a suite of functions for processing and visualizing Ribo-seq and RNA-seq data, facilitating ORF analysis within the gene structure or transcript context. All data files are imported and stored as R6 objects, leveraging object-oriented principles for efficient data handling. The package now supports a broader range of input formats, including bedGraph and bigWig, enhancing its compatibility with diverse sequencing datasets. The workflow and example code are available on the *ggRibo* GitHub page. The design, function, and basic steps for *ggRibo* visualization are summarized below.

### Annotation import

Annotation files are imported using the gtf_import() and eORF_import() functions.

- gtf_import(): Imports GTF/GFF annotation files, creating a Range_info object that extracts genomic features such as exons, transcripts, coding sequences (CDS), and untranslated regions (UTRs). This object serves as a foundational dataset for subsequent analyses.
- eORF_import(): Imports user-defined extra ORF (eORF) annotations, creating an eORF_Range_info object to accommodate non-standard ORFs not included in standard annotations.

### Data import

For data import, the *create_seq_input()* function is introduced alongside the existing *Ribo_data()* function.

- *create_seq_input()*: Constructs input lists for RNA-seq and Ribo-seq data by automatically detecting file formats (BAM, bigWig, bedGraph) and organizing the data into a structured format for downstream analysis. It supports both paired-end and single-end RNA-seq data and accommodates various Ribo-seq file types, enhancing the package’s flexibility.
- *Ribo_data()*: Reads Ribo-seq data files (or other single-nucleotide resolution data) and organizes them into a list of data frames, each containing read counts, chromosome numbers, positions, and strand information, enabling efficient manipulation and analysis.

### Visualization

The visualization capabilities are expanded with the following plotting functions:

- *ggRibo()*: The core plotting function, visualizing Ribo-seq periodicity alongside RNA-seq coverage within the gene structure context. It integrates transcript isoforms and genome annotations to highlight unannotated ORFs and unconventional translation events, now supporting bedGraph and bigWig input formats alongside BAM. It allows customization of aesthetics, scaling, and inclusion of additional sequencing data (e.g., TI-seq, degradome sequencing, CAGE-seq).
- *ggRNA()*: Plots RNA-seq data alone, offering visualization of transcription levels independent of Ribo-seq data.
- *ggRibo_decom()*: Generates a plot that decomposes Ribo-seq signals into three separate plots, one for each reading frame, providing insights into translation dynamics.
- *ggRibo_tx()*: Plots RNA-seq and Ribo-seq coverage in transcript coordinates, offering a detailed view of translation events at the transcript level by mapping data to spliced exon sequences.

### Basic steps for running *ggRibo*

1. *gtf_import*: Input genome annotation. Optional: use *eORF_import* to import the transcript and ORF ranges for extra ORFs recorded in gtf or gff3 format.
2. *create_seq_input*: Input RNA-seq files with *rna_files*, Ribo-seq files with *ribo_files*, sample names with *sample_names*. *ggRNA* only takes in RNA-seq files in this step.
3. *ggRibo*: Input a transcript id with *tx_id*. Alternatively, input a gene ID with *gene_id,* and the first transcript sorted by R will be used for *tx_id*. Users can input an eORF.tx_id that exists in the eORF gtf/gff3 file. Similar parameters work for *ggRNA*, *ggRibo_decom* and *ggRibo_tx*.

These functions utilize the *ggplot2* framework for high-quality, customizable plots, seamlessly integrating with the package’s object-oriented design. Two other sets of functions work behind the scenes for frame assignment and track plotting.

### Frame assignment

Several frame assignment functions are provided to assign translated reading frames accurately:

- *assign_frames()*: Assigns reading frames to Ribo-seq reads based on provided CDS ranges, enabling precise determination of translated codons.
- *assign_frames_extended()*: Extends annotated frame assignments into UTRs, accommodating overlapping eORFs for accurate frame calculations in complex regions.
- *assign_frames_with_extension()*: Allows custom extensions into the 5’ and 3’ UTRs (via fExtend and tExtend parameters), offering flexibility for analyzing alternative translation start sites or stop codon readthrough events.
- *exclude_eORF_reads()*: Filters out Ribo-seq reads overlapping with eORF regions from the main dataset, preventing redundancy in visualization.

### Functions for plotting the DNA sequence and gene model tracks

Additional functions enhance the visualization of DNA sequences and gene models:

- *plotDNAandAA()*: For genomic regions ≤ 201 nucleotides (via the plot_range parameter in *ggRibo(), ggRibo_decom()* and *ggRNA()*)), this function generates plots displaying DNA nucleotides and corresponding amino acids, aiding in codon usage and reading frame analysis. For regions > 201 nucleotides, it highlights start codons (green) and stop codons (red) in each frame.
- *plotGeneTxModel()*: Creates gene models illustrating exons, UTRs, CDS, and eORFs for specified genes and isoforms, adjusting for overlapping features and enhancing clarity with customized colors and labels.
- Two similar functions: *plotDNAandAA_tx()* and *plotGeneTxModel_tx()* perform identical functions as their parent functions for the *ggRibo_tx*() function.

The coding for the *ggRibo* package and parts of this Method section were assisted by the ChatGPT model o1 and Grok v3, with initial code adapted from a modified version of *RiboPlotR* (using base R plots) and transformed into *ggplot2* by AI tools and HLW.

## Supporting information

All supplemental material

## SOFTWARE AVAILABILITY

*ggRibo* is available on GitHub (https://github.com/hsinyenwu/ggRibo). The example data are stored on Mendeley Data: *Arabidopsis* (https://data.mendeley.com/datasets/wm6cS5zbtw/1) and human (https://data.mendeley.com/datasets/m3t293k4wr/1)

## ACKNOWLEDGEMENTS

This work used computational resources and services provided by the Institute for Cyber-Enabled Research at Michigan State University. This work was supported by a predoctoral training award under Grant Number T32-GM110523 from the National Institute of General Medical Sciences of the National Institutes of Health to IDK, and research grants from the National Science Foundation under Award Number 2425390 and the National Institute of General Medical Sciences of the National Institutes of Health under Award Number R35GM155375 to PYH. The content is solely the responsibility of the authors and does not necessarily represent the official views of the NSF or NIH.

## AUTHOR CONTRIBUTIONS

HLW developed the *ggRibo* package and tested it with *Arabidopsis* and tomato data. IDK tested *ggRibo* with human and rice data and fixed a few bugs. HLW and IDK prepared the GitHub tutorial. HLW and PYH wrote the paper with input from IDK.

