## Supplementary material for "A ggplot-based single-gene viewer reveals insights into the translatome and other nucleotide-resolution omics data": All supplemental material

Supplemental Figure S1. A summary of *ggRibo* improvements compared to *RiboPlotR*

Supplemental Figure S2: Comparing 'gene' view among *ggRibo*, *GWIPS-viz*, and *RiboCrypt*

Supplemental Figure S3. Comparing 'transcript' view between *ggRibo, Trips-viz,* and *RiboCrypt*

Supplemental Figure S4. Schematic output for *ggRNA*, *ggRibo_decom*, and *ggRibo_tx*

Supplemental Figure S5. Examples of *ggRibo* plots for the human *IFITM3* gene

Supplemental Figure S6. Examples of *ggRibo* and *ggRibo_tx* plots in rice

Supplemental Figure S7. Zoom in plots for *Arabidopsis TAS3* for the regions at the start and stop codons

Supplemental Figure S8. Plotting *Arabidopsis TAS3* gene using *ggRibo_decom* function

Supplemental Figure S9. Example of a novel sORF identified within an lncRNA using the 'blue' reading frame

Supplemental Figure S10. Decomposition of Ribo-seq reads for nested ORFs

Supplemental Figure S11. Comparing multiple Ribo-seq and RNA-seq datasets reveals differential isoform translation in *Arabidopsis* tissues

**Supplemental Figure S1. A summary of *ggRibo* improvements compared to *RiboPlotR.***

| | ***RiboPlotR*** | ***ggRibo*** |
|---|---|---|
| Plot system | Base R | ggplot2 |
| Object oriented programming | No | Yes (R6) |
| Annotation style | Transcript id = gene id + . + number | Any gtf/gff3 accepted by GenomicFeatures |
| Number of samples | 2 | Any as RAM allows |
| ORF types visualized | uORF, mORF | uORF, dORF, mORF ouORF, odORF, oORF, ncORF |
| Visualize Ribo-seq reads on ncRNA | No (need a defined ORF) | Yes |
| Zoom in on a specific region of a gene | No | Yes (use plot_range) |
| Adjust the y-axis of Ribo-seq to see lowly translated events | No | Yes (use Ribo_fix_height) |
| Extend main ORF frame to 5′ UTR or 3 ′ UTR (check non-AUG start or stop codon readthrought) | No | Yes (use fExtend for 5′ UTR and tExtend for 3′ UTR) |
| Change frame colors from red, blue, green | No | Yes (use frame_colors) |
| Plot DNA sequence | No | Yes (set show_seq = TRUE, and input FASTA file) |
| Plot direction | As genome browser | Always left to right (always 5′ -> 3′) |
| Accepted data formats | RNA-seq (bam file)<br>Ribo-seq (tabular file) | RNA-seq (bam file, bigWig, bedGraph)<br>Ribo-seq (tabular, bigWig, bedGraph) |

### Supplemental Figure S2

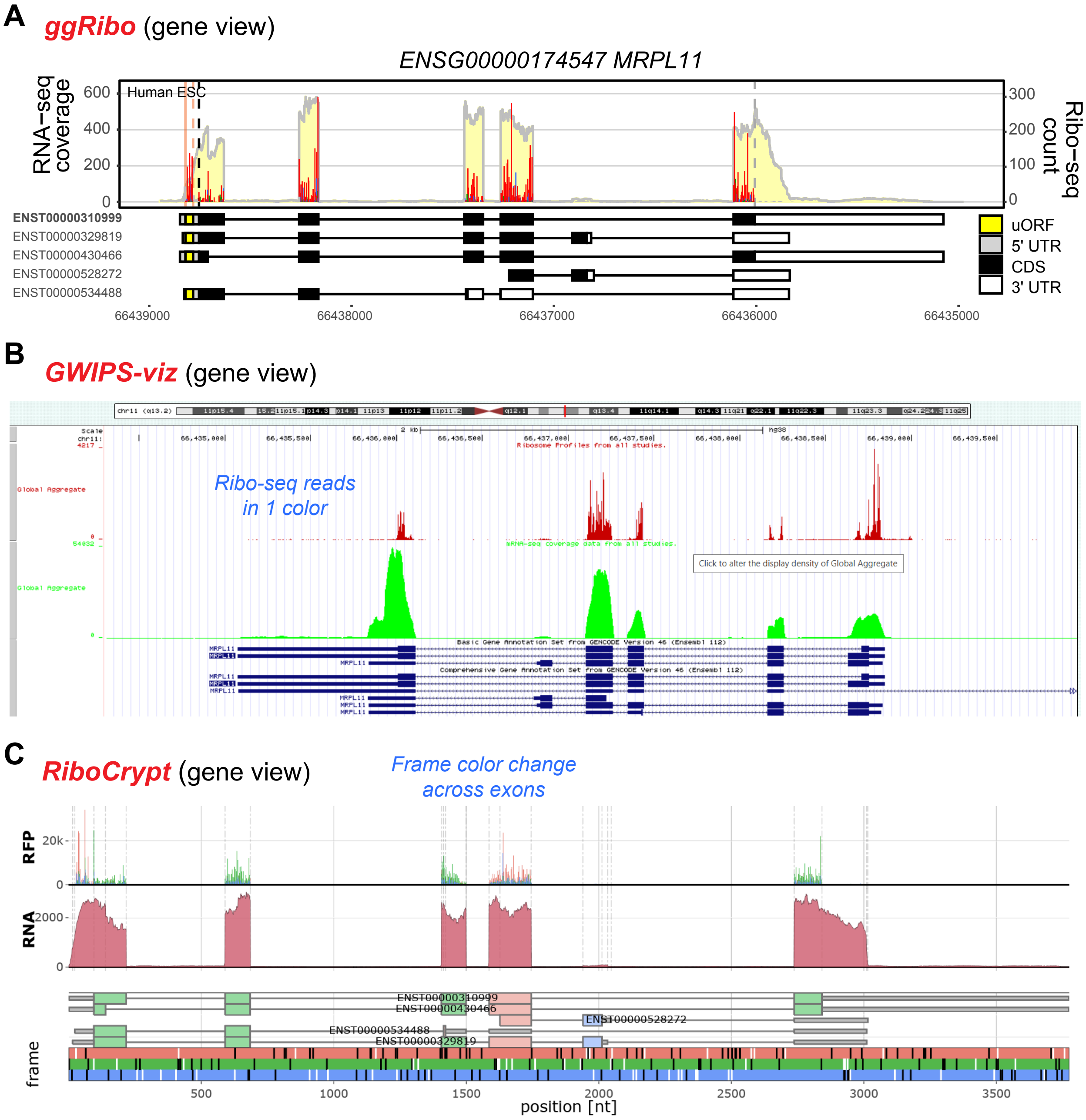


**Supplemental Figure S2: Comparing 'gene' view among *ggRibo*, *GWIPS-viz*, and *RiboCrypt.*** Ribo-seq/RNA-seq data for human *MRPL11* gene were plotted using *ggRibo* and two other tools (*GWIPS-viz* and *RiboCrypt*). Comments in blue italic fonts were added in Illustrator.

(A) *ggRibo* plots the Ribo-seq periodicity and RNA-seq coverage in the context of gene structure, while simultaneously integrating gene and isoform annotations. Major expressed isoform(s) can be visually identified by cross-referencing RNA coverage with isoform models. The Ribo-seq reads are color-coded by reading frames; enrichment of one color (here, red) indicates strong 3-nt periodicity for active translation. Notably, *ggRibo* assigns frame color based on the coding sequence of a transcript, ensuring consistent frame coloring across all exons for an ORF. This design is different from *RiboCrypt* (see below).

Note that unannotated ORF(s), such as a uORF here, can also be defined in the gene model. The Ribo-seq and RNA-seq data are from (Chothani et al. 2022).

(B) *GWIPS-viz* (https://gwips.ucc.ie/cgi-bin/hgGateway) displays Ribo-seq and RNA-seq data on separate plots and provides gene structure context. However, the Ribo-seq reads are shown with a single color, so reading frames cannot be distinguished. The data presented here is an aggregate of all human data collected on *GWIPS-viz* (Michel et al. 2018).

(C) *RiboCrypt* (https://ribocrypt.org) also displays Ribo-seq and RNA-seq data on separate plots and provides gene structure context in 'gene' view. However, *RiboCrypt* assigns frame colors based on the genomic DNA sequence. Thus, the frame color can change across different exons for an ORF. The data presented here is an aggregate of all human data collected on *RiboCrypt*.

### Supplemental Figure S3

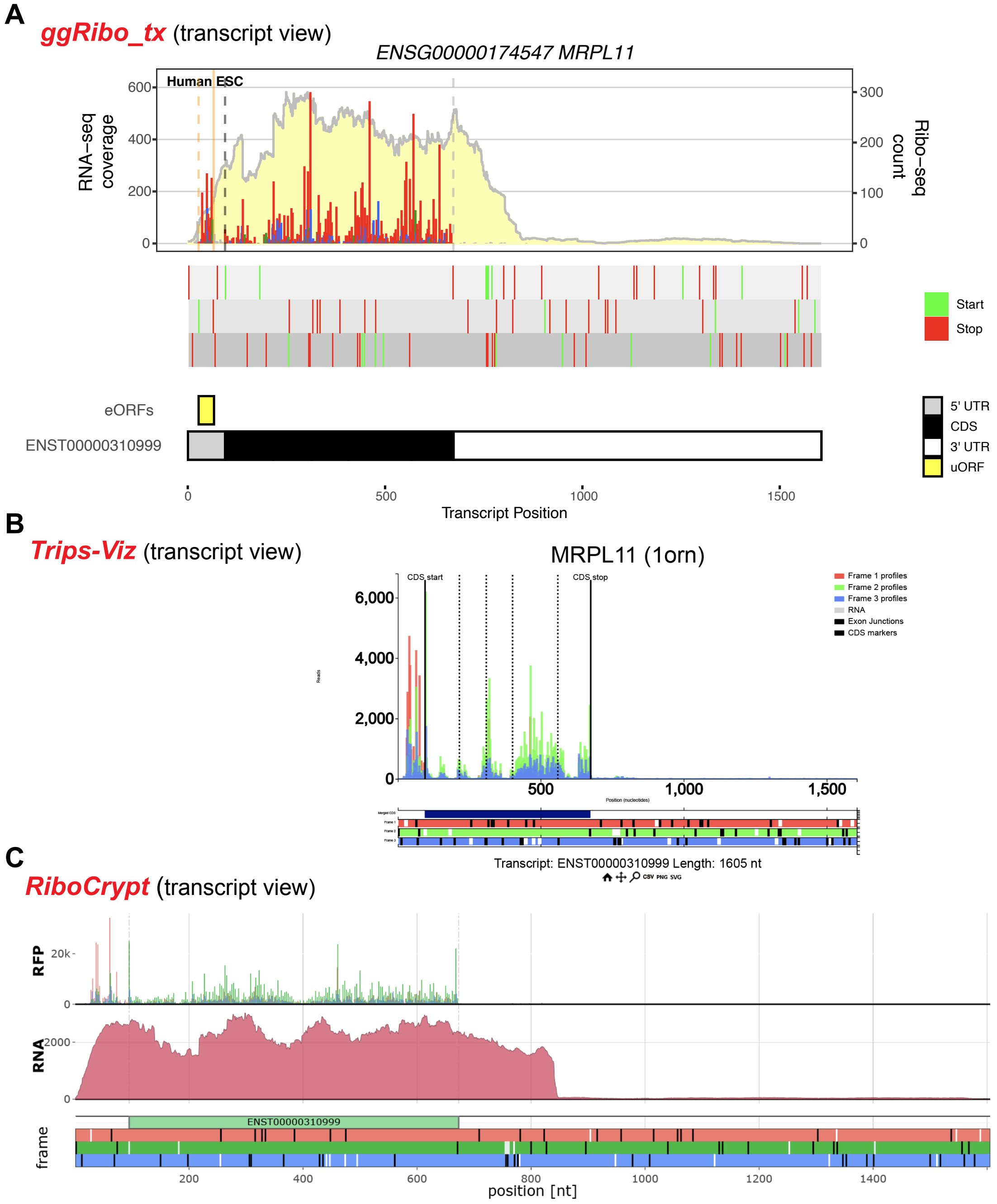


**Supplemental Figure S3. Comparing 'transcript' view among *ggRibo*, *GWIPS-viz*, and *RiboCrypt*.** Ribo-seq/RNA-seq data for the transcript ENST00000310999 of the human *MRPL11* gene were plotted using *ggRibo*, *Trips-Viz,* and *RiboCrypt.*

(A) Transcript view using *ggRibo_tx* function in *ggRibo*. An optional track highlighting start and stop codons in each frame is shown. The transcript model showing 5′ UTR, CDS, and 3′ UTR, as well as a uORF, is also displayed. The Ribo-seq and RNA-seq data are from (Chothani et al. 2022).

(B) Transcript view using *Trips-Viz*. The data presented here is an aggregate of all human data collected on *GWIPS-viz* (Michel et al. 2018).

(C) Transcript view using *RiboCrypt*. The data presented here is an aggregate of all human data collected on *RiboCrypt* (https://ribocrypt.org).

### Supplemental Figure S4

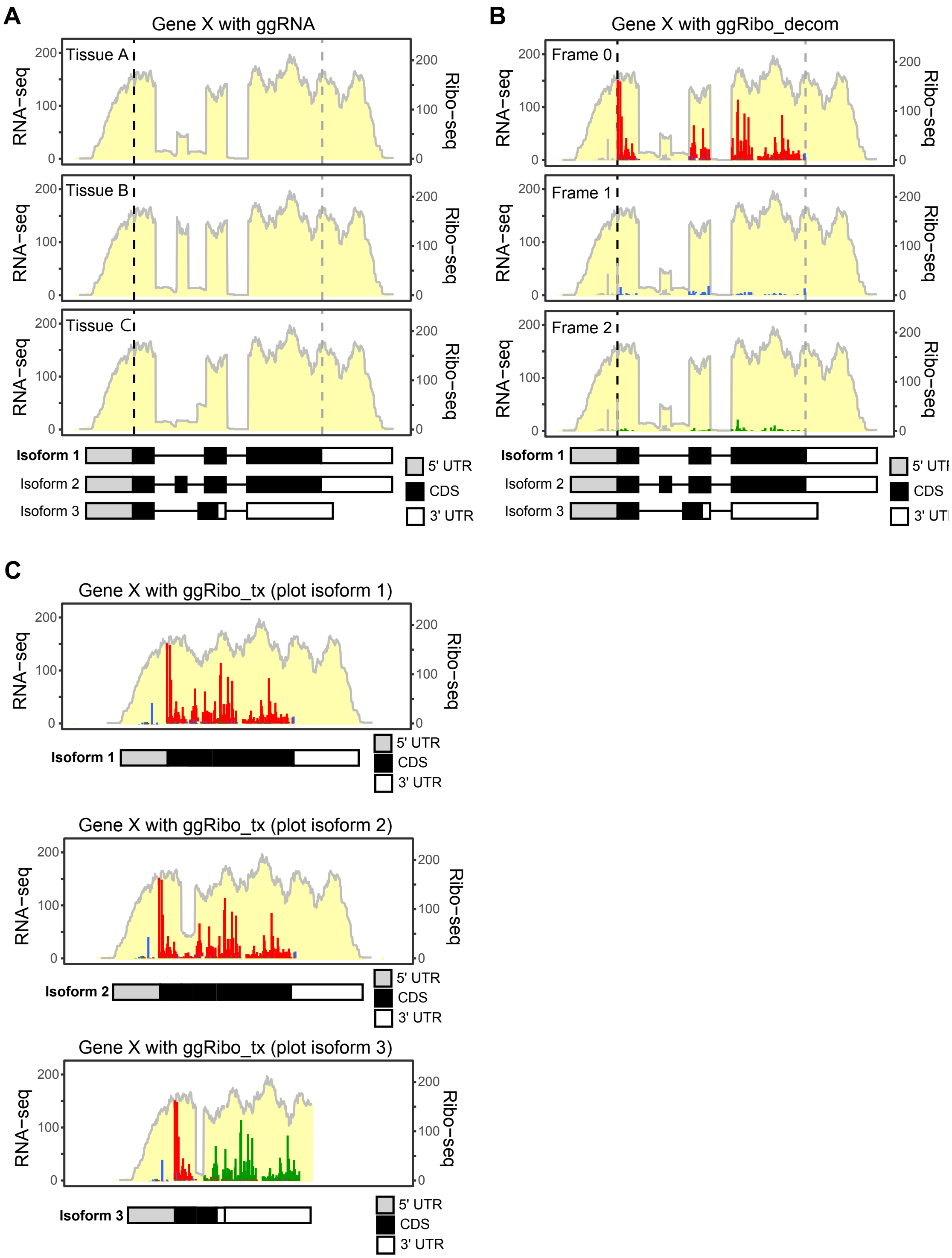


**Supplemental Figure S4. Schematic output for *ggRNA*, *ggRibo_decom*, and *ggRibo_tx*.**
(A) *ggRNA* plots the coverage of RNA-seq.
(B) *ggRibo_decom* plots Ribo-seq reads that are in different reading frames in separate plots. Users can use *ggRibo_decom* to visualize the read distribution in different frames.
(C) *ggRibo_tx* plots Ribo-seq and RNA-seq reads in transcript coordinates. Note that if a poorly expressed or unexpressed isoform is plotted (e.g., isoforms 2 and 3 in this example), a large intron-like gap or a region enriched for out-of-frame reads may appear.

**Supplemental Figure S5**

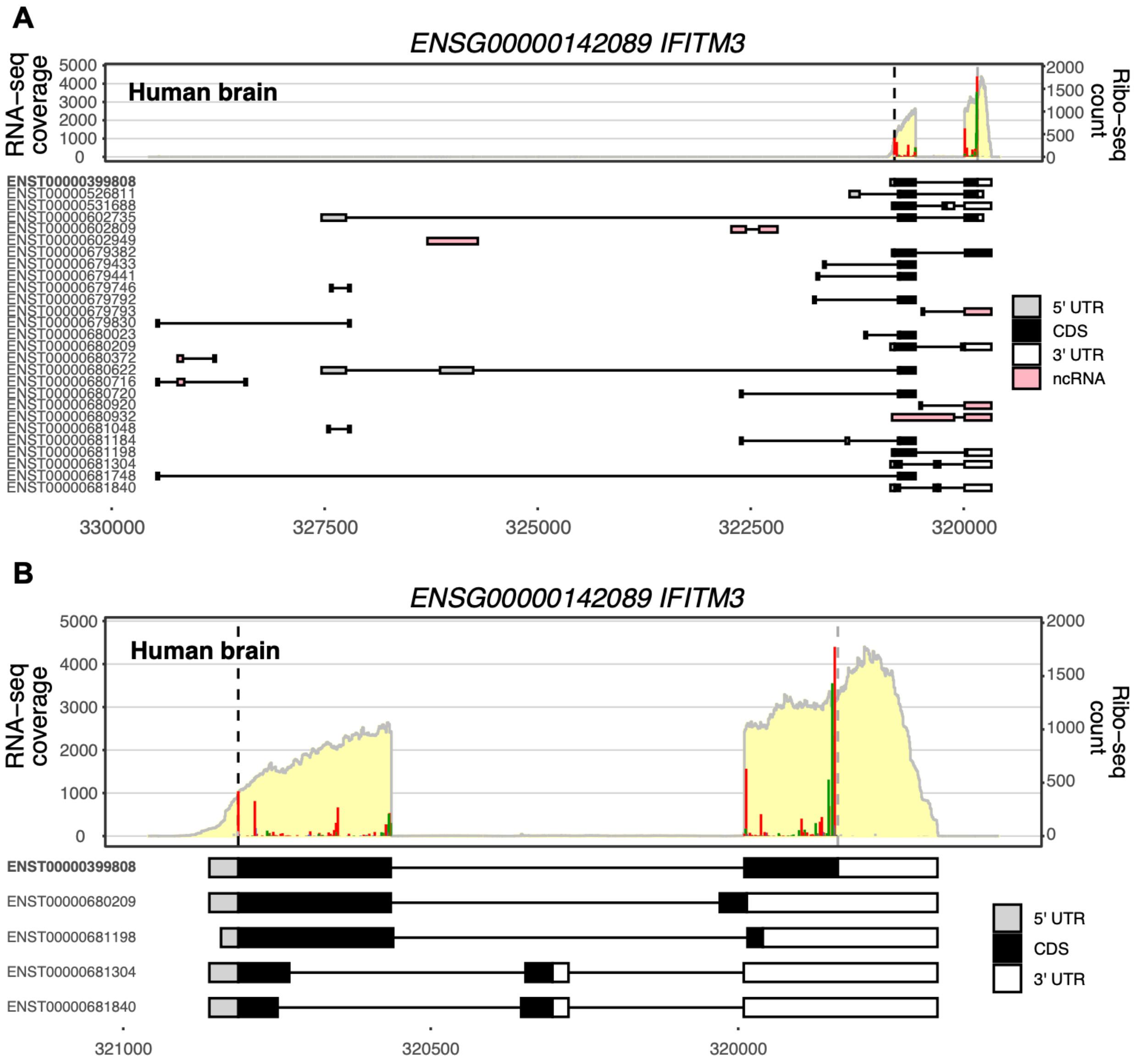


**Supplemental Figure S5. Examples of *ggRibo* plots for the human *IFITM3* gene.** Plotting RNA-seq and Ribo-seq profiles of the human *IFITM3* gene using *ggRibo*. The human brain Ribo-seq and RNA-seq data are from (Chothani et al. 2022).

(A) A gene view of the *IFITM3* gene. Although multiple isoforms are annotated (including some without an annotated CDS), it appears that only the ENST00000399808 isoform (bolded) is expressed and translated in this particular tissue.

(B) A closer view of ENST00000399808 and a subset of other isoforms in the same region confirms that ENST00000399808 is the major expressed and translated isoform.

### Supplemental Figure S6

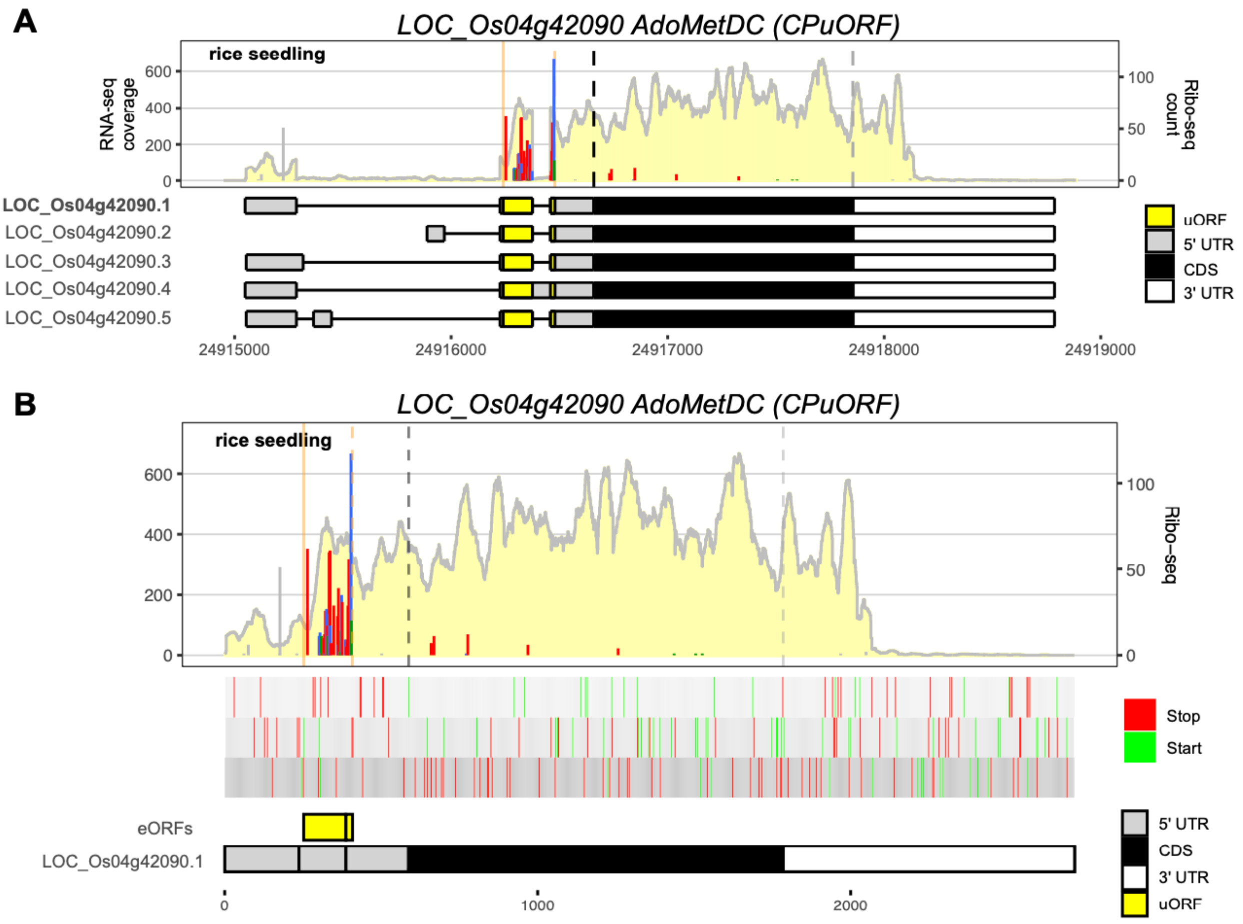


**Supplemental Figure S6. Examples of *ggRibo* and *ggRibo_tx* plots in rice**

Plotting the rice *AdoMetDC* (*AdenosylMethionine DeCarboxylase*) gene using *ggRibo* (A) and *ggRibo_tx* (B). *AdoMetDC* genes contain a conserved peptide uORF (CPuORF) that is conserved between *Arabidopsis* and rice (Franceschetti et al. 2001; Hayden and Jorgensen 2007). Ribo-seq and RNA-seq data are reported in (Yang et al. 2021).

**Supplemental Figure S7**

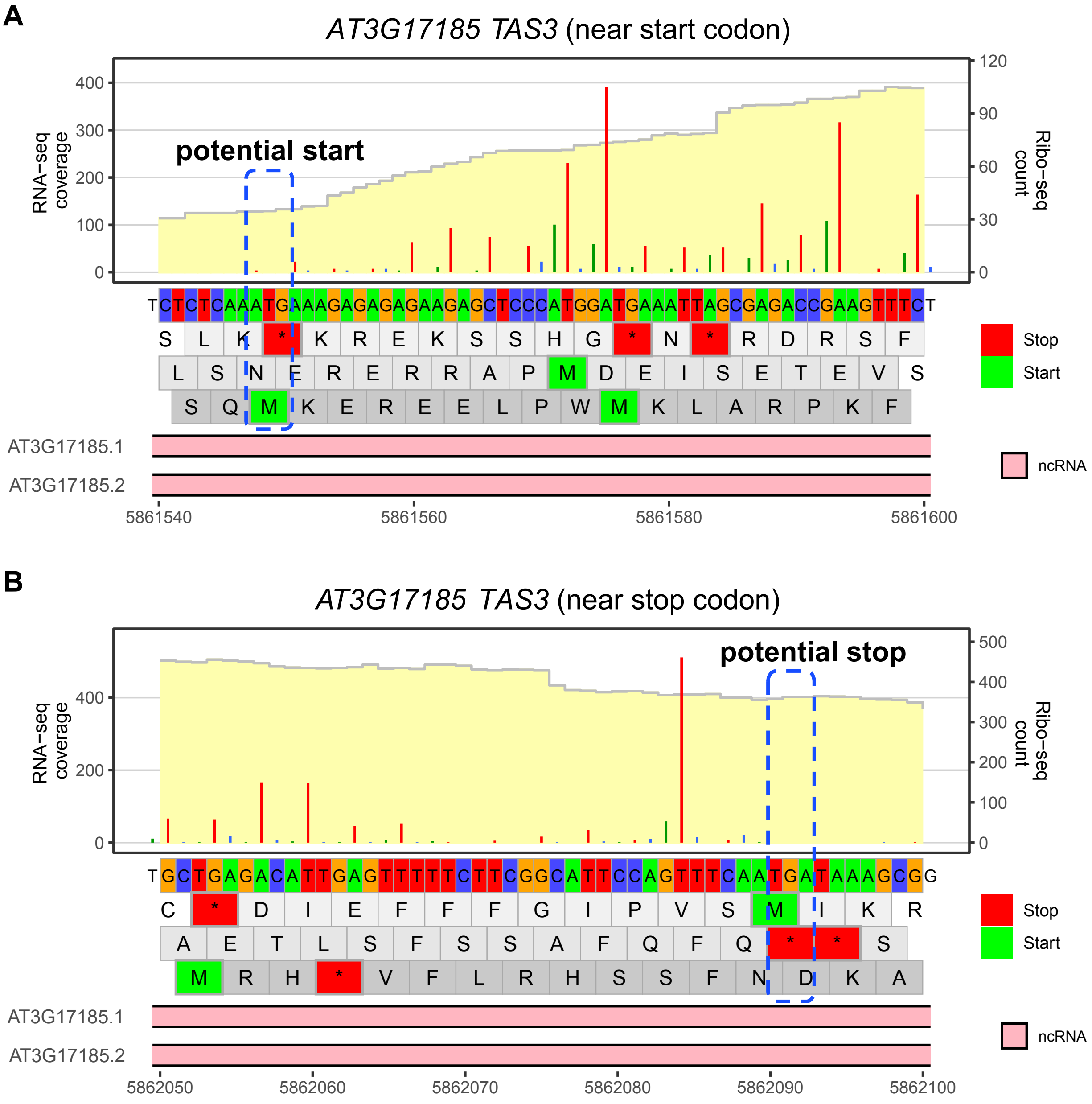


**Supplemental Figure S7. Zoom in plots for *Arabidopsis TAS3* for the regions at the start and stop codons**

Regions near the start codon (A) and stop codon (B) of *TAS3*. The potential start and stop are highlighted and annotated in Adobe Illustrator.

**Supplemental Figure S8**

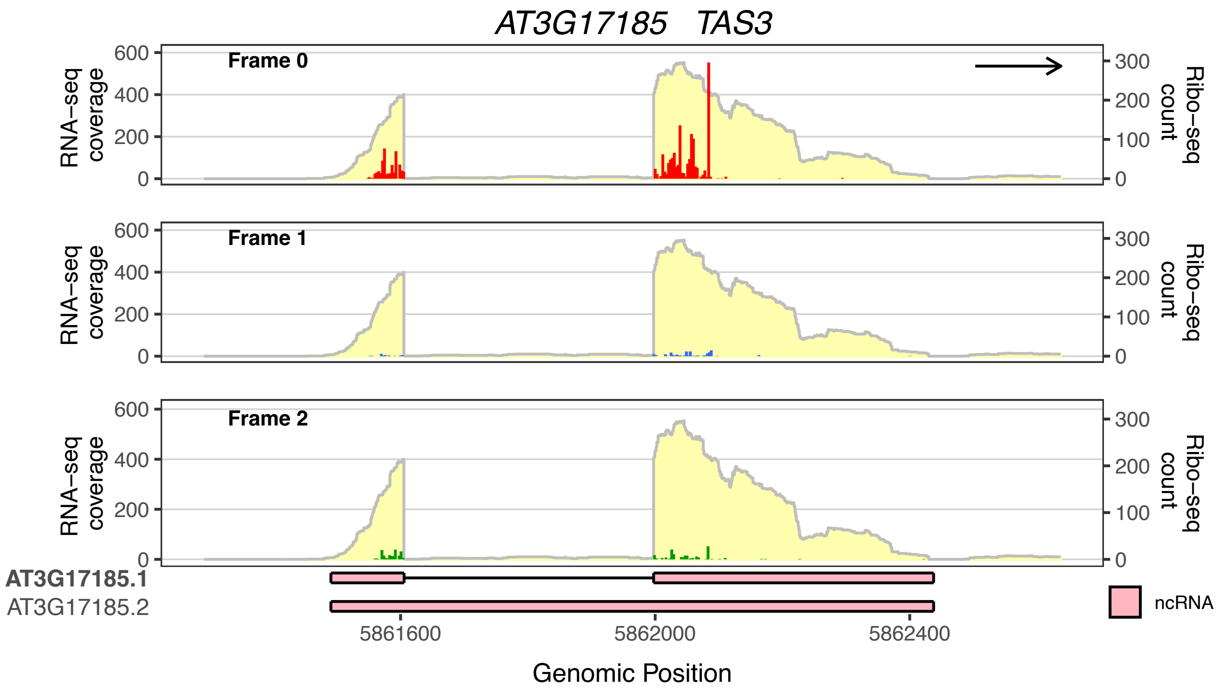


**Supplemental Figure S8. Plotting *Arabidopsis TAS3* gene using *ggRibo_decom* function.** *ggRibo_decom* function is used for the decomposition of the Ribo-seq reads in different reading frames for *TAS3*. The arrow on the top right (plotted with the option '*plot_genomic_direction*=TRUE'), which points to the right, indicates that the gene direction is consistent with the direction shown on the genome browser. Ribo-seq and RNA-seq data are reported in (Wu et al. 2024a).

**Supplemental Figure S9**

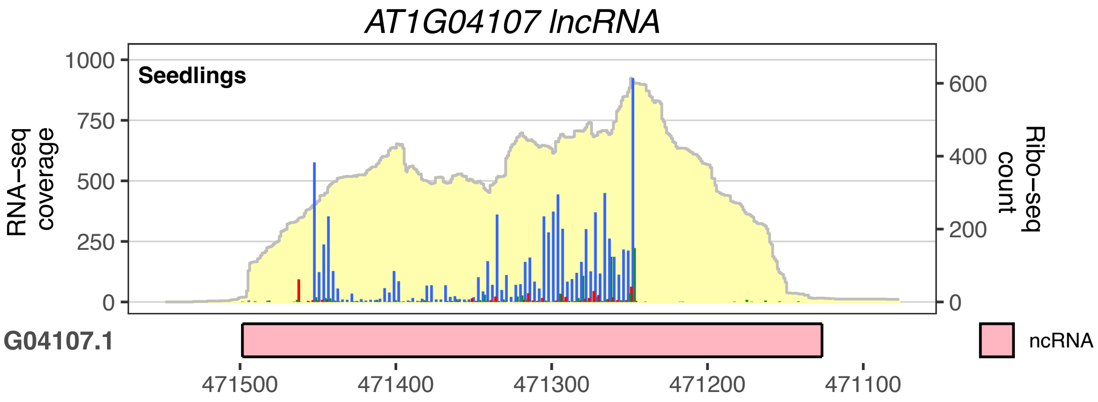


**Supplemental Figure S9. Example of a novel sORF identified within a lncRNA using the 'blue' reading frame.** Ribo-seq and RNA-seq expression profiles of a novel sORF identified in an annotated lncRNA in *Arabidopsis* seedlings (Wu et al. 2024a) are shown. The strong 3-nt periodicity of Ribo-seq reads in the 'blue' reading frame suggests the presence of a translated sORF.

### Supplemental Figure S10

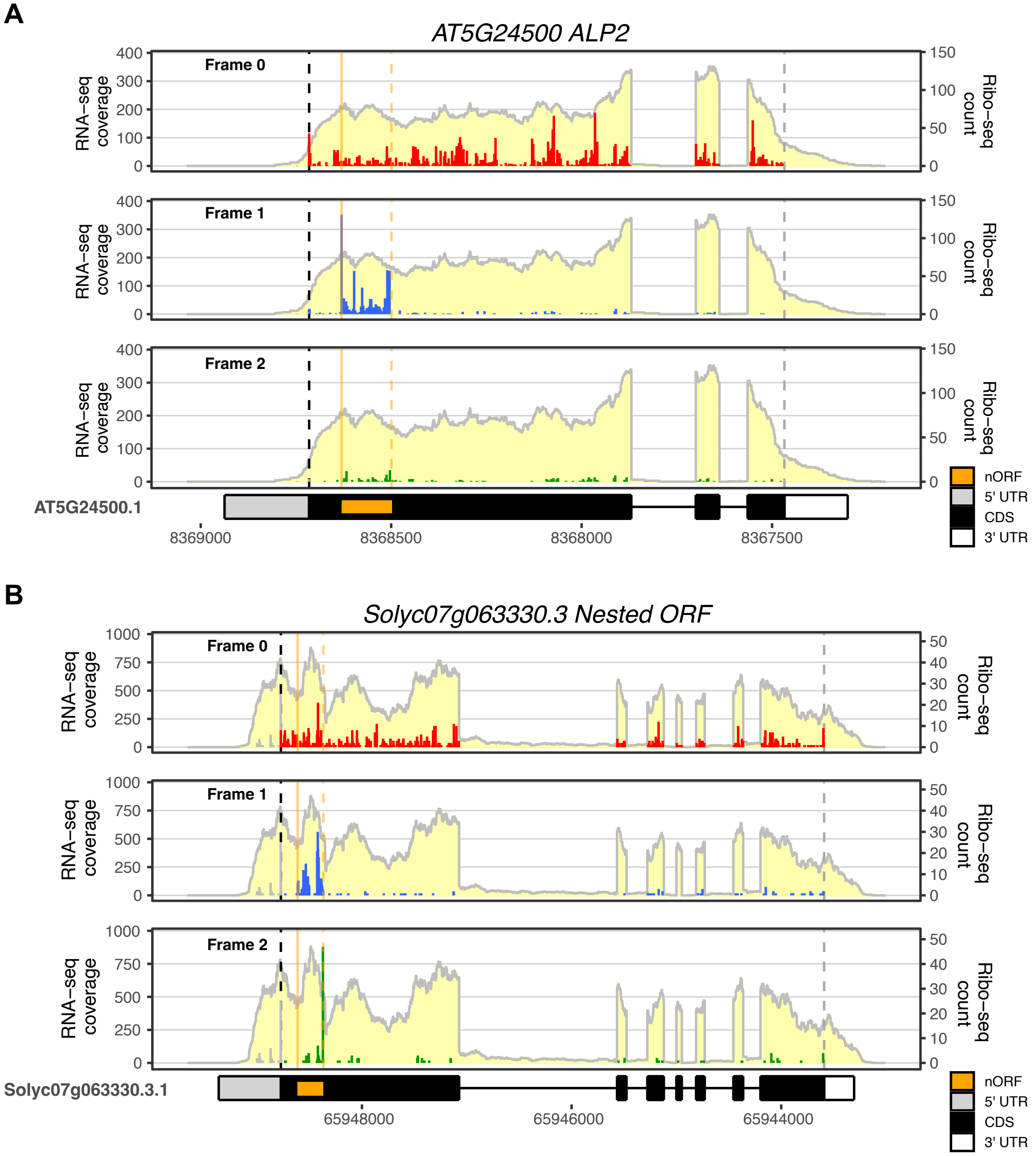


**Supplemental Figure S10. Decomposition of Ribo-seq reads for nested ORFs.** **(A, B)**: The two nested ORFs shown in Figures 4E and 4F are plotted by *ggRibo_decom* function to show the read distribution in each frame. In both cases, we can clearly observe the translation of the nested ORFs using the blue reading frame.

**Supplemental Figure S11**

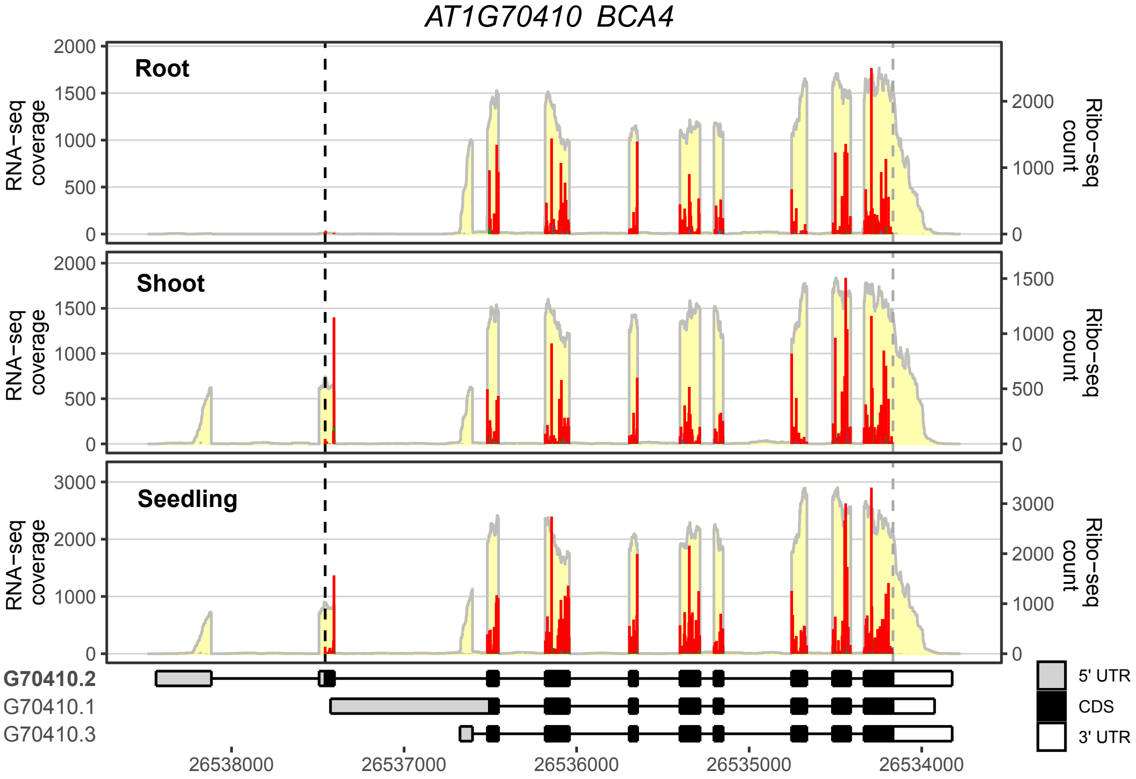


**Supplemental Figure S11. Comparing multiple Ribo-seq and RNA-seq datasets reveals differential isoform translation in *Arabidopsis* tissues**

Ribo-seq/RNA-seq expression profiles of *BCA4* in *Arabidopsis* root, shoot, and whole seedling are compared. The root and shoot data were reported in (Hsu et al. 2016), and the seedling data were reported in (Wu et al. 2024a). Note that isoform 2 is nearly undetectable in the root, but is highly expressed and translated in the shoot and whole seedling. Because the 5' UTR for isoform 3 was overestimated in the Araport11 annotation, its first exon was manually shortened to match the expressed exon range using Adobe Illustrator.
